# Geometrically encoded positioning of introns, intergenic segments, and exons in the human genome

**DOI:** 10.1101/2025.05.29.656862

**Authors:** Luay M Almassalha, Kyle L MacQuarrie, Marcelo Carignano, Cody Dunton, Ruyi Gong, Joe Ibarra, Lucas M Carter, Wing Shun Li, Rikkert Nap, Parambir S. Dulai, Igal Szleifer, Vadim Backman

## Abstract

Human tissues require a mechanism to generate durable, yet modifiable, transcriptional memories to sustain cell function across a lifetime. Previously, we demonstrated that nanoscale packing domains couple heterochromatin (cores) and euchromatin (outer zone) into unified reaction volumes that can generate transcriptional memory. In prior work, this framework demonstrated that RNA synthesis occurred within the ideal zone (intermediate density) portions of the domain. Naturally, this creates a question of where genes are positioned in relation to the packing domain architecture and which genetic material fills the domain core to sustain transcription. Here we propose that this could be solved by the encoded positioning of introns, intergenic segments, and exons as a projection of the functional packing layers of domains. This suggests that introns and intergenic segments are coupled to adjacent exons to generate coherent packing domain volumes. We illustrate how this organization would reconcile contradictions in epigenetic patterns, non-randomness in oncogenic mutations, and produce durable transcriptional memory. We conclude by showing that this genome geometry might have coincided with the rapid evolution of body-plan complexity, suggesting that chromatin geometry could be fundamental to metazoan evolution.

## Introduction

Although most cells in a multicellular organism share the same genome, they differentiate into hundreds of cell types. Even cells of the same lineage can perform different functions, shifting over timescales from minutes to decades, while many disease states arise from altered gene expression. A key underlying question is how genetic information gives rise to diverse and dynamic cell phenotypes across development, aging, and disease. Many sequence-dependent transcriptional regulatory elements are well studied, including enhancers, promoters, insulators, silencers, etc^[1–4]^. These provide significant insight into transcriptional regulation. Paired with imaging data, an emerging view is that chromatin organizes into packed structures throughout the nucleus. We recently described the structure-function lifecycle of nanoscale packing domains to result in a functional coupling between heterochromatin and euchromatin into a unified functional volume^[5,6]^. The formation and function of chromatin packing domains intersects physical and biological processes. This process is self-assembling, with transcriptional reactions having the capacity to initiate domain formation. As a result, it introduces a mechanism for transcriptionally mediated cellular memory encoded as a physical structure. An interesting physical feature of these packing domains is that density exists as a continuous gradient. This gradient, when paired with the size of enzymes (heterochromatin enzymes are small, ∼2nm compared to 6nm for euchromatin enzymes), generates a mechanism to geometrically guide enzymes to different locations within the volume^[6]^. Interestingly, an intermediate zone is generated that appears to be optimal for the positioning of RNA polymerase II (Pol-II) and transcription factors. Paired with modeling and super resolution imaging, we showed that this efficient packing – heterochromatin deposition into cores - acts to increase transcriptional output at distal regions^[6]^. This suggests that the genome is folded in a manner that can couple heterochromatin and euchromatin together into unified functional structures.

Applying this information to muscle differentiation, we observed that the activation of transcription on exons on the genes Myh1 and Myh2 was associated with the deposition of heterochromatin within non-exonic (NE) segments (introns, intergenic segments) of gene bodies^[6]^. Naturally, we wondered if this observation provides a system for genes to pack into nanoscale structures where NE elements act to produce the packing domain volumes that could position exons into the ideal zones. Using myogenesis again as a model system, to our surprise, we observed that 58 of the top 100 differentially activated genes in muscle differentiation are associated with decreased accessibility in NE segments (**Figure 1a-c**). This behavior contrasts with accessibility at the transcription start site, which is generally stable (**Figure 1b-c**). Given this finding, we then examined whether heterochromatin resides in NE segments with adjacent RNA synthesis across human tissues broadly by analyzing data available through ENCODE^[7–9]^. In multiple tissues, critical functional genes contain heterochromatin within NE segments adjacent to actively transcribed exons (**Figure 1d**). For example, H3K9me3 (a marker of constitutive heterochromatin) is deposited within TNNI3 (cardiac troponin) in cardiac samples. This biallelically expressed gene must be transcribed throughout the human lifespan to maintain heart contractility (**SI Figure 1**)^[10–13]^.

**Figure 1.**
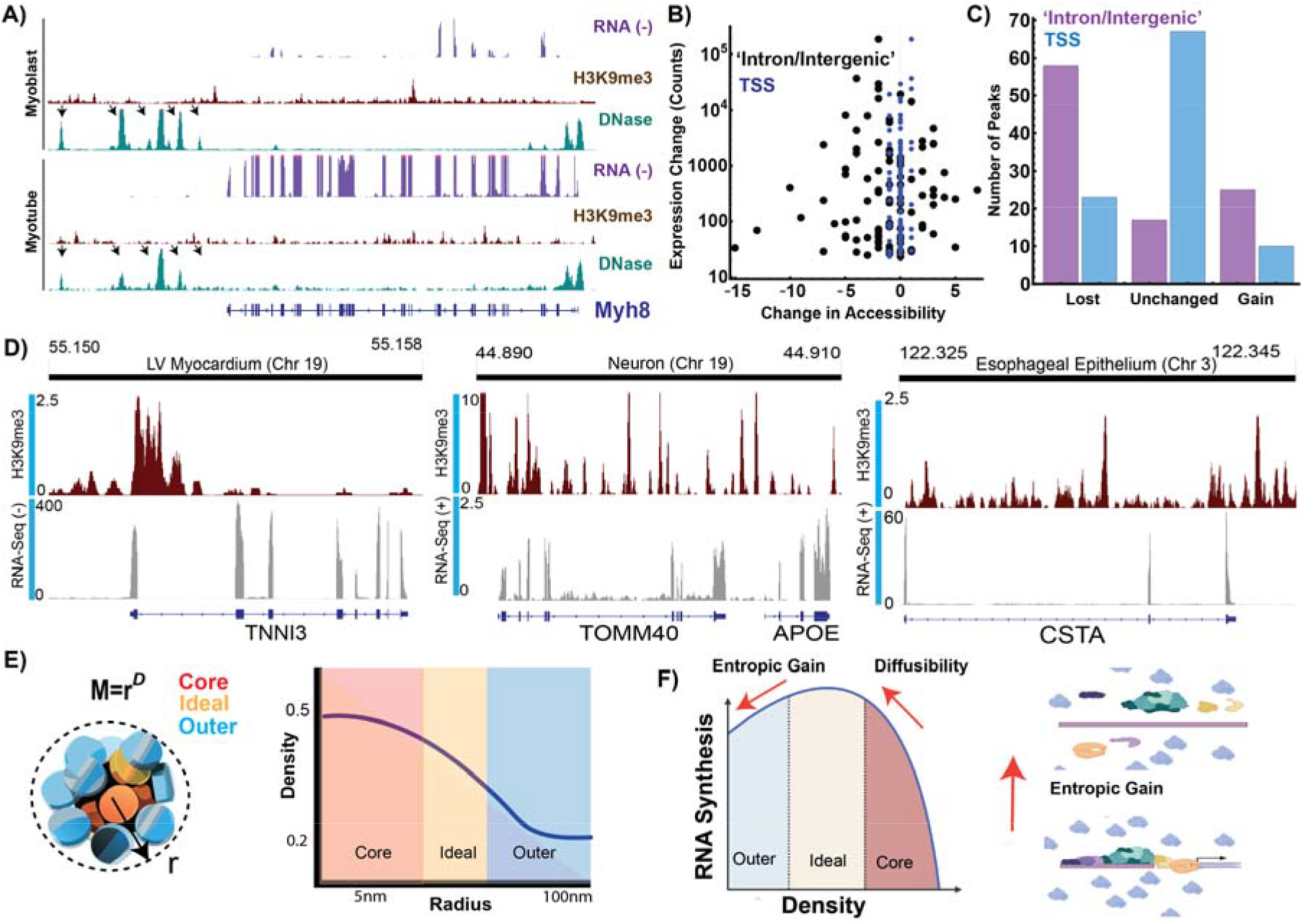
A paradox of the loss of accessibility with transcriptional amplification. **A)** DNAse-seq analysis demonstrates loss of accessibility with concurrent transcriptional activation during muscle differentiation. **B-C**) Loss of accessibility in non-exonic segments occurs across myogenic transcriptional activation. **D**) Transcriptional activity is associated with intronic heterochromatin deposition across tissue types. These are biallelic, constitutively expressed genes in their respective tissues. Critically, this includes TNNI3 (troponin), which is a component of the sarcomere necessary for cardiac contractility. **E**) Mass-fractal packing domains create a unified reaction-volume due to the continuous density gradient across domain layers. **F**) Transcription is non-monotonically dependent on local density due to the trade-off between entropic gain (remaining bound as an intermediate complex decreases the excluded volume to other macromolecules) and the diffusibility of reactant species. As density continues to increase, Pol II subunits can no longer penetrate deeper into a domain volume resulting in inhibition.

We wish to propose a radical hypothesis – described within this work – that nanoscale packing geometry is encoded in the positioning of exons, introns, and intergenic segments as projections of the functional layers of domains observed on ChromSTEM tomography. This would pair NE segments with adjacent exons as a linear projection of the 3-D volumes observed experimentally^[6,14–16]^. For this hypothesis to be valid, it requires (1) a length-dependent pairing of NE segments with exons to generate volumetric ratios, and (2) that they be oriented in the direction of gene transcription. We demonstrate that both properties are observed. We then provide experimental evidence from nascent-RNA sequencing, ChIP-Seq, ATAC-Seq, paired-end tag sequencing (ChiaPET), and ChromSTEM tomography to support this hypothesis. While the proposed geometric system produces a mechanism to generate transcriptional efficiency, memory, and complexity, we show that it comes at increased mutation risk of the exposed branch points between domains. By analyzing oncogenes, we show mutation frequencies across all cancers correlating with genes containing branch point segments. Of particular interest is that this geometric system may have emerged in parallel with increased body plan complexity of metazoans, suggesting that geometry introduces a possible novel, non-mutagenic system to increase information sampling based on physical properties. In sum, this theory generates new avenues of research across diseases of aging, species evolution, and complex organ development rooted in physical genomics.

## Results

### Chromatin domain geometry optimizes transcription in the human nucleus

All methods encounter limitations in measuring chromatin at the smallest length-scales due to the problem of missing information. Many tools provide sequence-dependent chromatin interactions including proximity capture methods (Hi-C, Sprite, etc)^[17–19]^ and *in situ* hybridization (FISH) imaging^[20–22]^. However, connectivity in a population may not equate to geometry in a single cell (**SI Figure 1**). To complement connectivity measurements with nanoscale density, most studies employ super-resolution imaging, accessibility assays, or variants of chromatin immunoprecipitation^*[14,15,23–30]*^. For example, due to probe size, it appears that antibodies used for super-resolution imaging of chromatin cannot penetrate high density domains *in situ*. This results in high-density and low-density regions devoid of signal (**SI Figure 2a-d**)^[31,32]^. A separate problem is present for super-resolution imaging using FISH probes that rely on formamide dehybridization. The denaturation process swells nanoscale domains, causing a loss of volume and density information ^[33–35]^. Due to these limitations of nanoscale methods, there is limited knowledge of human gene geometry at the smallest length scales. While FISH has shown that a large chain or melt phenotypes are present for a few very genes, the observed distances are consistent with these genes being compressed in space (**SI Materials and Methods**)^[36]^. With the advent of ChromEM/ChromSTEM imaging, it is now possible to study how chromatin transitions from disordered beads-on-a-string into 3-D volumes as a general framework, but this too lacks sequence-specific information ^[15,16]^. To overcome this limitation, it is necessary to pair the findings from this modality with other imaging modalities, molecular technologies, and modeling. From this integrated approach, it was shown that chromatin domains observed on ChromSTEM imaging appear to be self-assembling structures guided by transcription and cohesin to create volumes. The geometry of these packing domains is a mass fractal: for large genes and loops, the mass of a gene/loop (in nucleosomes) fills space as a function of radius by the dimension, *D*. This is analytically described by the equation, *Mass*(*nucleosomes*) = *r*^*D*^; a property confirmed on ChromSTEM tomography^[15,16,37,38]^.

In many ways, it is conceptually easier to consider two limiting cases of mass fractals to understand the structure of nanoscopic packing domains. A random walk without attractive potentials or confinement has scaling *D*=2, which has a uniform distribution of density within a volume (**SI Figure 3a**). An extension of this is a confined random walk, the fractal globule, which will have a contact scaling of *D=3* but also has a uniform distribution of density (**SI Figure 3b**)^[38]^. These two extremes are similar in that density is unform – they resemble statistically homogeneous structures that are similar to spheres with different levels of packing. It may be tempting to assign chromatin as an open state to *D=2* and closed state to *D*=3^[38]^. However, this is not observed experimentally on ChromSTEM. Instead, chromatin throughout the nucleus organizes into domains with *D* values ranging between 2.2 and 2.8 with a radius ranging between 25-175nm. This indicates non-uniform density: a gradient is present that radially decays from a high-density interior to a low-density periphery ^[15,16]^. The result is that packing at the nanoscale couples high density (heterochromatin cores) with a lower density outer zone (euchromatin). An intermediate density between these two regions appears to be an ideal zone for transcriptional reactions to occur. Based on these zones, it was shown that disruption of heterochromatin enzymes can experimentally decrease RNA synthesis due to the disruption of the ideal zone supported by the core element. Furthermore, it appears that this system can self-assemble through transcription-mediated loops first generating nascent (small, poorly packed) domains. While these results indicate that transcription appears to happen at the intermediate region, it remains to be understood where the geometric information for such a system is stored.

A possibility we consider is that large genes and loops are partially constrained (packed) into domain volumes. We can consider muscle and myosin heavy chain 1 (Myh1) as a representative example for comparison between models. Myh1 is ∼26,000 basepairs, which translates into ∼130 nucleosomes. A nucleosome is approximately a cylinder composed of ∼200bp with diameter of ∼11nm and height of ∼5.5nm. Stacking end-to-end to produce a fully stretched state produces a ∼715nm loop. For scale, chromosome 2 is composed of 243Mbp, contains ∼1,200 genes, and in mature muscle cells has a radius of ∼1.5*μ*m (**SI Figure 4)**^[39]^. Since muscle function requires the synthesis of hundreds of genes, many of which are quite large, accounting for space is crucial. Outside of muscle, a similar problem arises with loops. Many loops are over 100kbp (which translates into 5.5um), a length which would span the radius of the nucleus if not folded in space (**Figure 3a, SI Figure 5**)^[40,41]^. We propose a hypothesis that packing information is stored by exons non-randomly pairing with NE in order to generate the functional layers observed on ChromSTEM imaging (**Figure 2b**).

**Figure 2.**
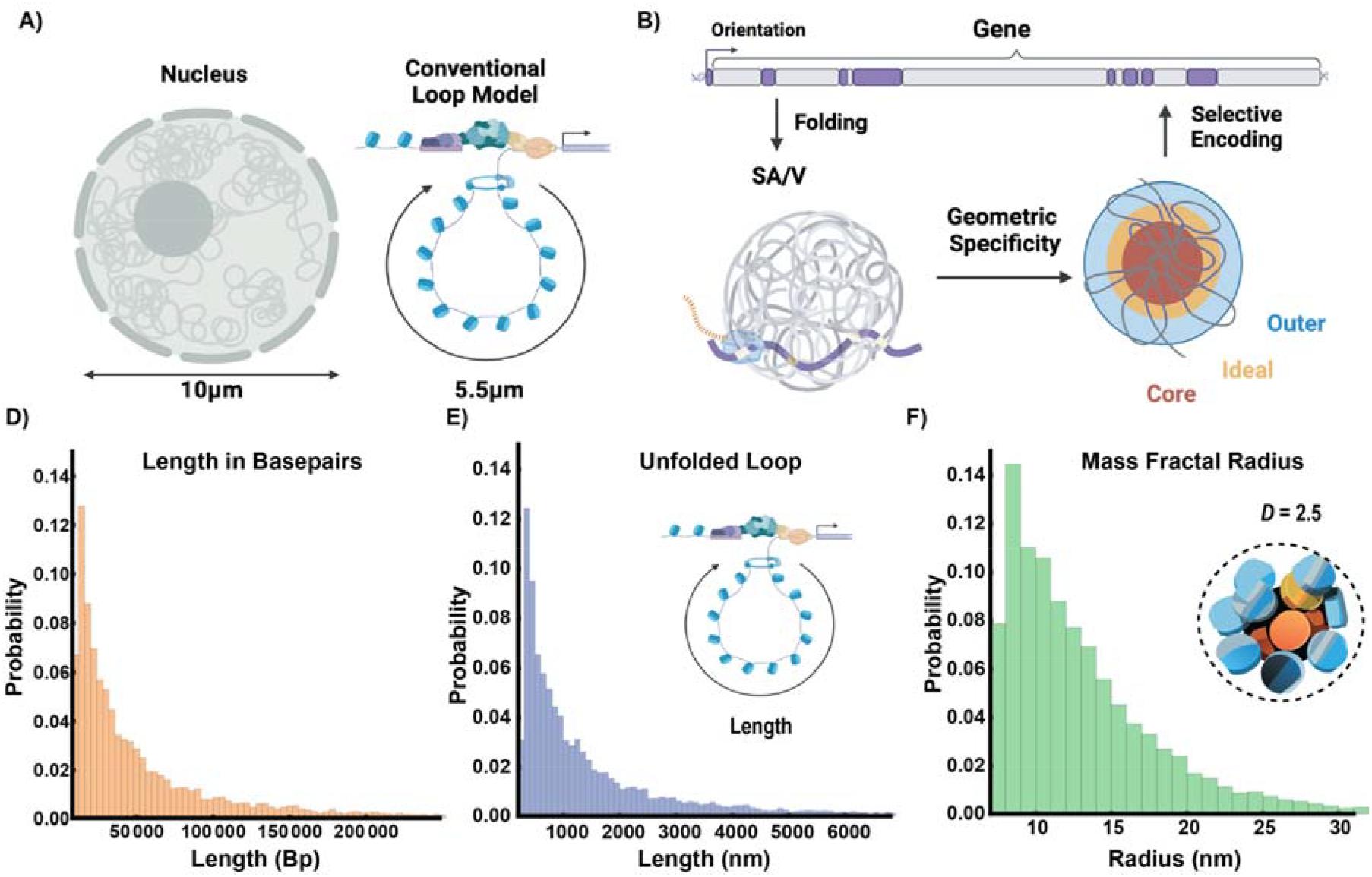
Packing domains as a geometric solution to optimize nuclear volume and transcriptional efficiency. **A)** Loops have a broad range of sizes with many larger loops (>100Kbp) generating lengths that would span the human nucleus without a system for efficient packing. **B)** Proposed framework that the position of exons, introns, and intergenic elements produces a system to reliably generate reaction volumes. Exons with short intronic sequences fold into an ideal zone within a volume generated by NE DNA (the ideal zone as a surface-area to volume - SA/V – of the total volume). The resulting volumes represent the structures observed on ChromSTEM imaging. The continued selection for elements across broad-timescales results in an encoding within the genome. **C-E**) Transformation from beads on a string into mass-fractal volumes compresses genes from micron-length chains into nanoscopic volumes.

**Figure 3.**
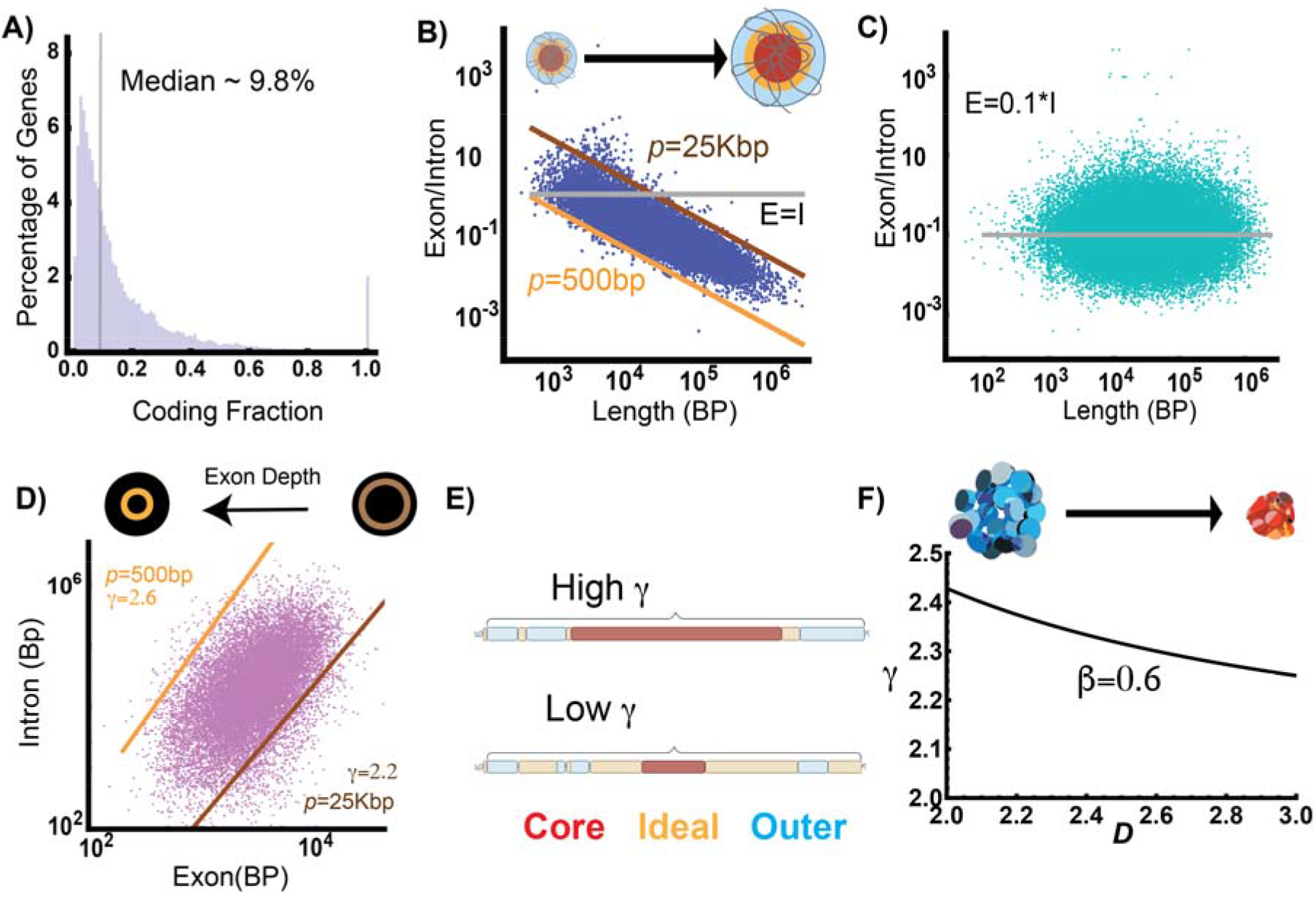
Introns and genes are geometrically linked to exon length by physical principles. **A)** Histogram of protein coding genes in the human genome showing that the median fraction is ∼9.8% exonic with a subset of genes that are almost completely exon. **B)** Plot of the ratio of exon (ideal zone)/intron (total volume) compared to the gene length of human genes. The constant, *p*, reflects the position along a domain volume. Here, *n* set to 1. As length increases, as expected, the volume ratio decreases as expected for a distribution of reaction volumes. **C)** Randomization of exon/intron segments results in the statistically grounded null hypothesis of no relationship between length and exon/intron ratios with a fraction approaching the median of 0.1. **D**) Intron length is a power-law of exon length for protein coding genes with values of *p* and *γ* as reported. **E**) Schematic representation of gene composition in relation to *p* and *γ*, indicating that high *γ* indicates more non-exonic volumetric elements are present within a segment. **F)** Relationship between *γ* and *D* depends on the proportion of the exons making up the ideal zone where *β*=1 indicates the entire exon contents are confined to a hard surface.

### Human genes are a predictable, power-law geometric assembly of exons and introns

As a consistent convention across the different types of RNA products, we refer to DNA transcribed and processed into functional RNA as ‘exons’ independent of the RNA product class (mRNA, lncRNA, pseudogenes, etc). Similarly, we refer to infrequently transcribed (NE) DNA as introns (within a gene body) and intergenic segments (between bodies)^[42–44]^. We recognize that non-exonic DNA has many sequence-specific regulatory functions^[45–47]^. We intentionally do not make sequence -specific measurements, as the focus is on packing geometry encoding novel information. Instead, this investigation focuses on whether 3-D geometry, guided by transcription, could project these elements as reaction volumes within the human genome. This geometric theory is as follows.

The proposed volumetric domain functional organization results in an inverse relationship of the ratio of exons-to-introns as a function of gene length. This is because the length of a segment (gene/loop) folded into a domain translates into the radius as a function of *D* (**Figure 2d-f**) ^[15,16,30,31,48–50]^. The inverse relationship of a reaction zone to the domain volume mirrors that of the surface-area-to-volume ratio observed in spheres (**SI Figure 6**), which decreases as a function of the radius (**Figure 2b, SI Materials and Methods**). However, instead of the case of each layer being a thin surface, it instead refers to content organized into zones within a total volume. In this case, it indicates that genes contain almost enough NE DNA to fold into domains to position exons for efficient transcription. In sharp contrast to this inverse relationship, chromatin organizing into beads-on-a-string would result in a linear relationship between the exonic fraction to the NE length (**SI Figure 6**). This is because as the number of nucleosomes increases, the amount of accessible DNA increases proportionally (**SI Figure 6**). The null hypothesis is that no volume-producing geometric relationship exists and elements within the genome are randomly spaced. This would result in genes having a random amount and positioning of intronic DNA as a function of length. Based on this, one would reasonably assume that exon length would be ∼10% of any gene body independent of the length. This is because only 10% of the gene length on average is exonic. Instead, there is a broad distribution in the exon fraction within genes (**Figure 3a**, interquartile range of 4.6% to 21.1%) that is length dependent, as we describe below.

Using the publicly available human reference genome GRCh38, with annotations from RefSeq, we tested if the inverse relationship occurs across the human genome^[51]^. We calculated the ratio of exons to introns vs total length for all protein-coding genes, finding that most human genes are described by this geometric principle (**Figure 3b, Materials and Methods**). This relationship is defined as follows by Equation (1) where *p* is a scalar in basepairs:

1) 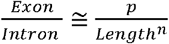

We observe most genes are well described by a *p* range from 500bp to 25,000bp, which is consistent with ∼2 to 125 nucleosomes when *n* is close to 1. This is observed both for individual isoforms or all the variant isoforms within the RefSeq database (**SI Figure 7**). An alternative explanation for such a trend is that as gene length increases, the exonic fraction decreases. If this were the case, this pattern would be observed even if exons are randomly redistributed. However, randomization produces a length-independent constant ratio of E/I of ∼0.1, indicating that gene length is non-randomly associated with exon content (**Figure 3c**). However, to account for the possibility that the observed ratio decreases due to length alone, we tested if intron length is geometrically a power-law of exon length.

If intron content is a power law with a value between 2 and 3, it would support the hypothesis that compositions of genes are related to packing geometry. Supporting our volumetric hypothesis, we observed this relationship (**Figure 3d**). This is captured by Equation (2) where *p* ranges from 500bp to 25,000bp:

2) *Intron ∝ Exon γ/p*

Understanding the function of the scalar, *p*, and the exponent, *γ*, requires considering how these translate linear information into 3-D volumes. Along the linear genome, *p* and *γ* are proposed to control the density of exonic information. Here, a lower *γ* (∼2) produces less spacing than larger *γ* (∼3). Likewise, the value of *p* indicates where along the depth of a volume a segment is being positioned (a smaller value suggesting it is deeper in a volume, and a larger value the inverse).

The central hypothesis would suggest that both degrees of freedom could interact to accommodate an effective packing configuration across multiple conditions (concentration of nucleosome remodeling enzymes, ion concentrations, nuclear size, existing domains, etc) that are difficult to measure in every human cell. Generally, a high *p* and low *γ* state produces high information density. As a result, structural genes such as Myh1 (*p* ∼7,800, *γ* = 2.2) are proposed to adopt more complex geometries with some degree of packing present. It is worth noting that in the proposed hypothesis, *γ* is inversely proportional to *D* as the transformation of the chain into the occupied volume, quantified by:

3) 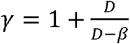 where β defines the fractional dimension of exonic contents on the ideal zone (**Figure 3e-f Materials and Methods**).

The values of these scaling exponents are indicative of whether exonic and NE regions likely follow 3D volume relationships of domains. In a case when the 3D packing geometry is not observed (e.g. a chain assembly), *β*=0 resulting in γ= 1, *c*= 1, and *n* = 0 with I ∝*E*: and *E* ∝ *L*. On the other hand, a strong packing geometry corresponds to 2 ≤ *γ* ≤3, *c*< 1, and n ∼ 1. Our experimental data is consistent with the latter case of packing geometry as we describe in detail within this manuscript. Since we propose that these properties are related to the activity of transcription, we verified that this behavior is independent of transcript type by analyzing the behavior of non-protein coding RNA classes (**SI Figure 7**).

### Gene positions on human chromosomes reflect the projection of 3-D domain volumes

We next investigated if these principles generalize to the positioning of genes on chromosomes. Since transcription guides domain formation experimentally, it suggests that genes oriented in *cis* may share domain volumes. In this hypothesis, intergenic segments could therefore share similar volumetric properties as introns to support domain formation. On both ChromSTEM imaging and in polymer simulations, packing domains are separated by an interdomain space formed by DNA ^[15,16,30,48,52,53]^. From experimental observations, this linker segment must contain an element that can recognize Pol II (for domain generation) and contain enough DNA to generate separation between the two domains. We refer to this as a ‘hinge’: when engaged by actively transcribing Pol II it will form separate segments into two domain volumes. A domain will be formed by the span between the two hinges. When not engaged by Pol II, a hinge may revert to fold into a volume element in an alternate configuration (**Figure 4a**). If hinges are only composed of transcribed segments, the supercoiling generated by polymerase could potentially alter the composition of the domains (domains could merge or separate randomly)^[54,55]^. Alternatively, if hinges were completely non-exonic, they would lack a mechanism for Pol II to guide domain segmentation. Since hinges are not domains, they are likely no more than a few nucleosomes long (<1000bp, ∼50nm long). Above this range, they would potentially interact with nucleosome remodelers via the mechanisms that produce domain volumes, limiting their ability to achieve spacing.

**Figure 4.**
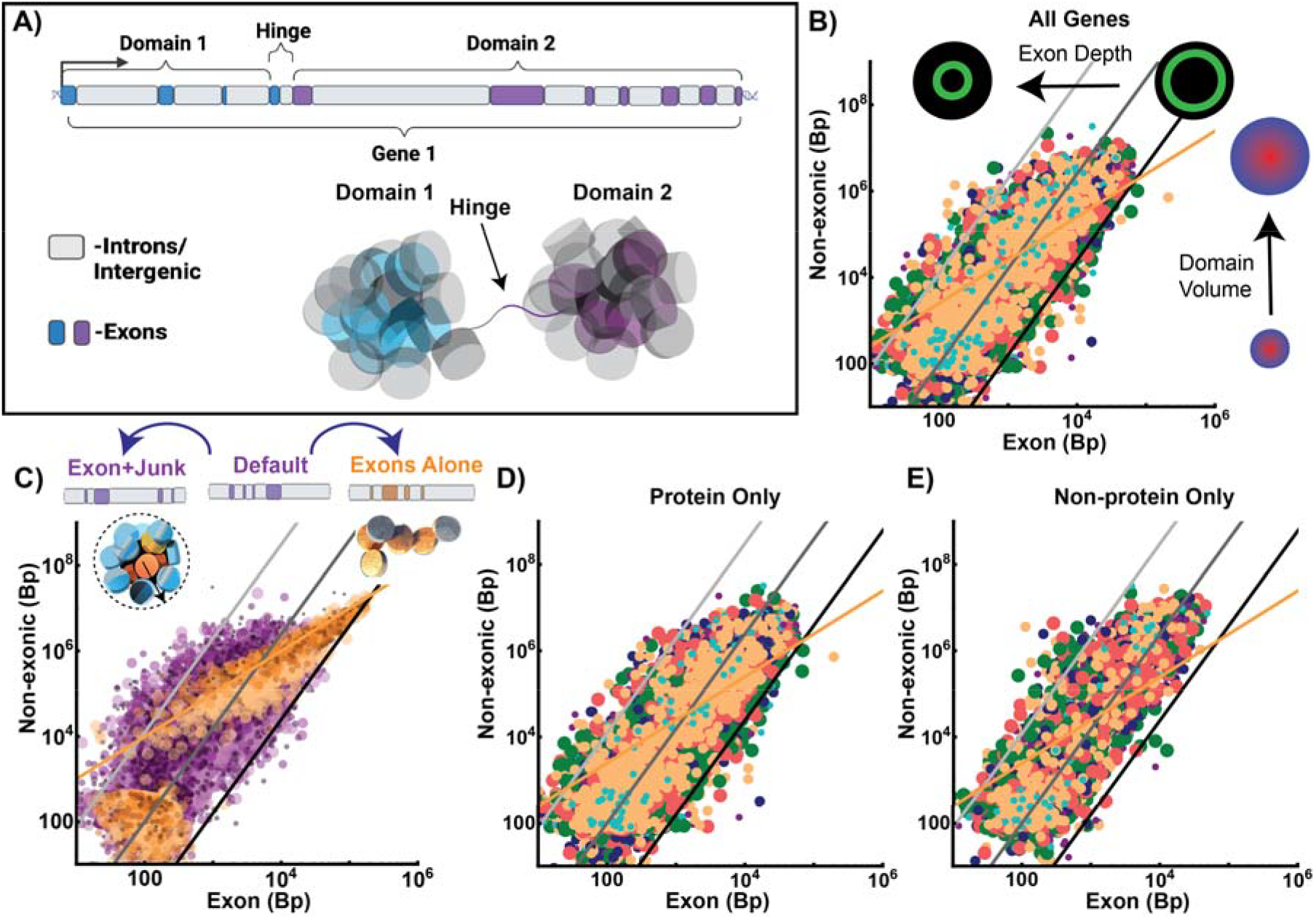
Exons are non-randomly coupled to adjacent volumetric DNA to generate power-law segments independent of the final RNA product. **A)** Schematic representation of the organization of a gene on a chromosome segmented into two separate domains generated by a hinge element. The reaction volumes are produced by the volumetric DNA to guide the position of exons to ideal reaction zones. **B)** Analysis of the structure of human chromosomes in the positive strand orientation showing power-law assemblies of nontranscribed (volumetric) elements scaling as a power-law of exonic (ideal zone) elements. **C)** Randomly redistributing an exon with adjacent volumetric DNA conserves power-law distribution. In contrast, randomly distributing exons results in a linear distribution of segments. **D-E**) Comparison of organization generated by considering only (**D**) protein coding genes compared to (**E**) only non-protein coding genes in the positive strand orientation. In either case, chromosomes assemble into power-law units.

By considering the presence of hinges, we find that every human chromosome assembles into power-law packing ratios (**Figure 4b**). This organization resembles individual genes described by Eq. 3, with *p* and *γ* adopting the same meaning:

4) *NE* ∝ Exon γ/*p*

This suggests that exons are non-randomly linked with their adjacent NE neighbor to produce reaction volumes. Since this is difficult to test experimentally, we analytically tested this hypothesis by randomizing the positions of exons alone and compared to randomizing exon+NE pairs (**Figure 4c**). If exons are uncorrelated NE elements producing packing domain volumes, then their randomization would still produce ratios that are consistent with volumes. When exons are distributed alone, the power-law compositions degrade into linear (unpacked) ratios (**Figure 4c-d)**. Conversely, when exon+NE are randomly distributed together, power-law patterns are maintained. This supports the hypothesis that exonic segments are non-randomly paired to an adjacent NE segment in 3-D volumes. If this is intrinsically driven by transcription itself and not the final product, this pattern would be present in either protein coding or non-protein-coding segmentation. Independent of the type of RNA generated (protein coding RNA, non-protein coding RNA), chromosomes assemble into power-law segments of exonic-/NE- elements (**Figure 4c-d)**. Collectively, these findings are consistent with the hypothesis that exons, introns, and intergenic segments are coupled in transcriptionally mediated reaction volumes.

### Predictions of genome geometry are observed across methods

The central hypothesis proposed in this manuscript – exons, introns, and intergenic segments are non-randomly coupled by the 3-D volume geometry – introduces four dependent and testable hypotheses, even as testing the central hypothesis directly is challenging:

H1) The DNA content of predicted segments resembles experimental observations on ChromSTEM,

H2)Exons have a higher likelihood of interacting with Pol II based on their position,

H3) RNA synthesis decreases from the introns of long genes from suboptimal positioning, and

H4) Hinge segments are enriched for features of active transcription.

Although these hypotheses cannot definitively prove the novel positioning hypothesis, as it would require manipulation and synthesis of genetic elements, the proposed central hypothesis requires all of them to be sustained. Hence we tested whether these dependent hypotheses are supported by experimental data.

To test H1, we utilized existing ChromSTEM imaging data from HCT-116 cells. The DNA content observed on ChromSTEM domains can be calculated by converting intensity into content using the mass-fractal relationship^[5,16]^. Although we cannot identify the sequence composition on ChromSTEM, we can compare the distribution of sizes between theory and experimental measurements of PDs. Consistent with theory predictions, the DNA totals generated by the segmentation described is very similar to those experimentally observed (**Figure 5a**, Mann Whitney U-Test 0.89 for negative reading frame and 0.93 for positive reading frame) but not for segments generated from randomly positioned exons (**Figure SI 8**, Mann Whitney U-Test <10^−45^).

**Figure 5.**
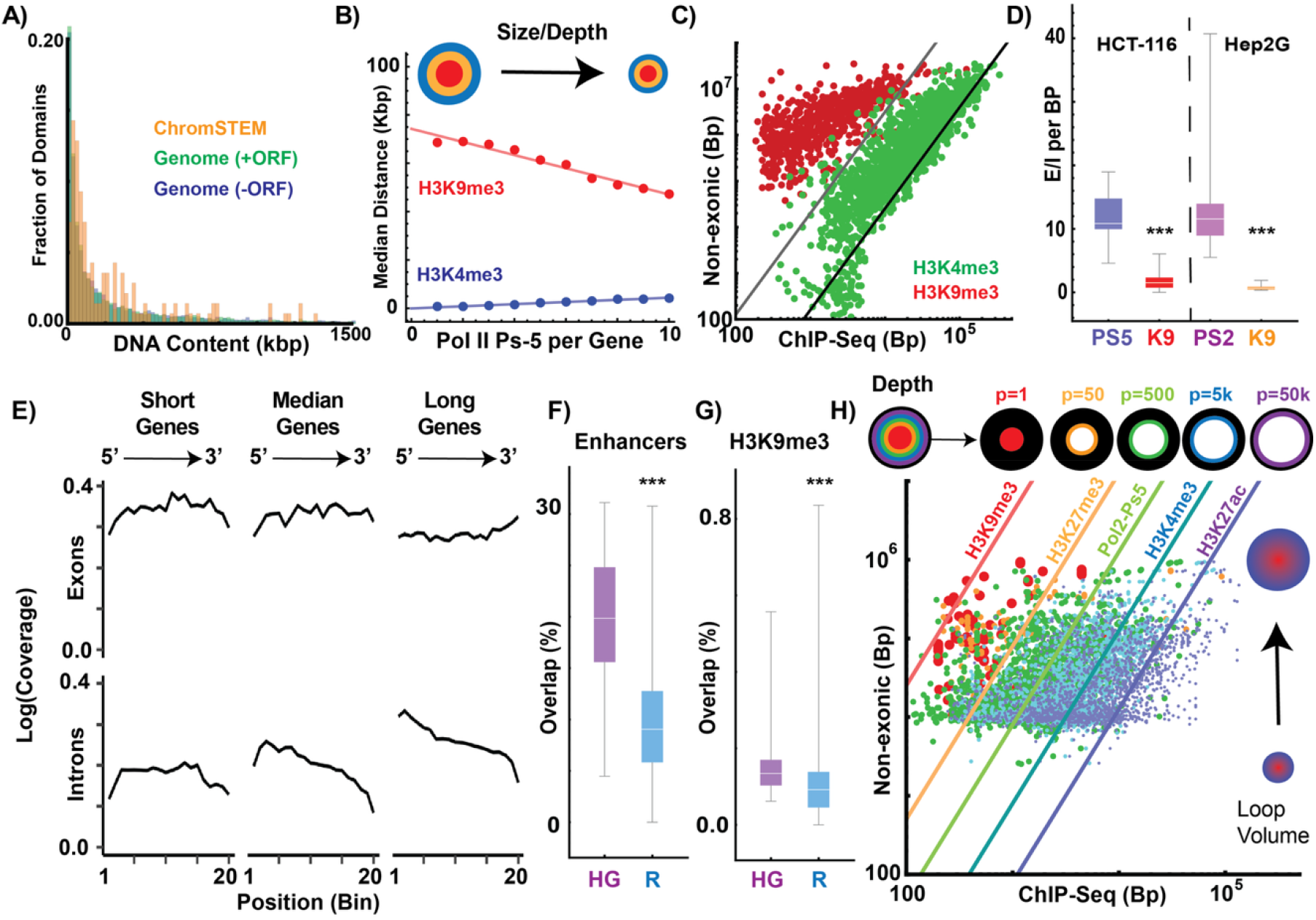
Exons, introns, and intergenic segments are non-randomly positioned into reaction volumes. **A)** Comparison of the DNA content observed in ChromSTEM packing domains compared to analytical predictions of the model when hinge length is less than 300bp demonstrating similar distributions in content. **B)** Analysis of distance between active Pol II (PS-5) to the nearest core element (H3K9me3) as a function of Pol II Ps-5 density. Shallower depths and smaller domains would result in a higher concentration of Pol II Ps-5 per segment and would be expected to have a shorter distance to a core as is experimentally observed. **C)** Analysis of the ChIP-Seq content of heterochromatin (H3K9me3) and promoter associated euchromatin (H3K4me3) within theorized volumes from the model in HCT-116. Consistent with underlying theory, chromatin segments are composed of both elements with positioning H3K4me3 coinciding with position of exon elements while heterochromatin is positioned to deeper layers. This partitioning suggests that heterochromatin is coupled with euchromatin in genetic segments. **D)** ChIP-Seq analysis of active isoforms Pol II Ps-5 in HCT-116 cells and Pol II Ps-2 in Hep2G cells within gene bodies (exons and introns) demonstrating preferential localization of polymerase onto exons per basepair length. If polymerases were uniformly throughout a gene body equivalent coverage per basepair would be observed. As a control for read-coverage bias, this was compared to H3K9me3 (K9) which preferentially localizes to introns. **E)** Nascent RNA-seq analysis in HCT-116 demonstrates nearly uniform synthesis of short introns and exonic sequences independent of gene lengths. In contrast, longer genes demonstrate decreasing synthesis of RNA in the direction of the reading frame consistent with Pol II having ideal reaction positions. **F**) Analysis of hinge segments in HCT-116 demonstrates an enrichment toward transcriptionally active features and enhancer positions compared to randomly generated segments (R). **G)** In contrast, a very small percentage (<1%) of hinge positions overlaps with constitutive heterochromatin. **H)** Experimentally observed RNA polymerase loops plotted as a function of their NE content (approximately the volume generated) compared to the observed ChIP-Seq content within each loop in HCT-116 cells. Within large polymerase loop domains, we observe an accumulation of heterochromatin. The total heterochromatin content increases a function of the size suggesting packing occurring as a function of the volume generated. Small loops are primarily composed of euchromatin, indicating their volume would appear to be relatively decompacted. Collectively, this suggests that transcriptional loops have a degree of packing that correlates to the packing behavior for domain volumes observed on ChromSTEM imaging.

H2 requires three observations. First, there must be a length-dependent coupling between functional zones (cores, active Pol II, and euchromatin) reflecting domain volumes. Specifically, the higher volumetric ratio in small domains produces a higher concentration of Pol II with a shorter linear distance to a core. In contrast to packing, a chain assembly would result in increasing distance between Pol II to H3K9me3. Using ChIP-Seq data available through ENCODE^[7–9]^, we tested if the coupling between transcription and heterochromatin is consistent with domain volumes by measuring the distance between active Pol II (Ps2/Ps5) and the nearest H3K9me3 as a function of the polymerase concentration^[6]^. We performed this analysis on induced pluripotent stem cells (GM23338), HepG2 hepatocellular carcinoma cells, and SK-N-SH neuroblastoma cells using data from ENCODE. If Pol II is guided by geometric position, an inverse distance relationship is likely to be conserved across models (higher Pol II concentration resulting in shorter distances). Further, because iPSCs are highly enriched in euchromatin ^[56,57]^, the theory predicts them to have the smallest domains with highest volumetric ratios and shortest distances. These predictions are observed: all three cell lines have the inverse distance relationship and iPSCs have the shortest distances on average (**Figure 5b, SI Figure 8**). Second, H2 indicates that exons are positioned to geometrically interact with polymerase, euchromatin, and heterochromatin. Specifically, one would expect H3K4me3 and active RNA polymerase II to localize in the projection of the ideal zone mainly composed of exons whereas H3K9me3 would localize to deeper layers. This would be manifested in the segments by a shift in the *p* required to position these elements such that H3K4me3 and Pol II would range between ∼250-25,000 bp and H3K9me3 to values below this range (1-50bp). This is indeed observed experimentally in HCT-116 cells (**Figure 5c, SI Figure 9**).

Third, H2 requires a Pol II preference to binding exons when accounting for the differences in segment length. This is similarly involved in H3 because if Pol II continuously moves throughout a gene, including long introns, it would not be uniformly distributed. In support of H2 and H3, active Pol-II is nearly ten times more likely to bind to an exon than an intron in both Hep2G and HCT-116 cells when accounting for length (**Figure 5d**, p-value <10^−3^). H3 further requires that this bias in positioning result in a gradual decay in the amount of RNA observed from introns in long genes due to decreased efficiency. In bulk RNA-seq, this results in a ‘saw-tooth pattern’ which was previously theorized to occur from the delayed processing of RNA by the splicing enzymes or the decay in introns^[58]^. However, the rate of splicing can greatly exceed the rate of transcription for long introns, indicating that this is not necessarily a rate-limiting event in RNA synthesis^[59–61]^. If genes are synthesized in total, the decay in intronic RNA likely would be symmetrical with the highest rates of decay in the shortest introns. This is because processing will not impact the synthesis rate within intron bodies and short segments will likely have the fastest processing. Utilizing nascent RNA-seq, we tested which configuration is most likely. Consistent with H3, there is a length-dependent decrease in RNA synthesis in 5’ to 3’ orientation for introns of long genes (**Figure 5e**).

We used a similar approach to investigate H4. We analyzed data from ENCODE and the Atlas of Enhancers^[2]^ in HCT-116 cells for hinges positioned in the human genome (HG) vs hinges generated by randomly repositioning exons (R). Hinge positions behaved as hypothesized, with enrichment in euchromatin marks and enhancers and depletion of heterochromatin markers compared to the random genome. In the human genome, ∼7.8% (interquartile range 7.1% to 9.5%) were bound by active RNA polymerase II (Pol II-Ps5), 9.7% (interquartile range 8.3% to 10.3%) were marked by H3K27ac, 8.7% (interquartile range 7.8% to 9.5%) were marked by H3K4me3, and 19.8% of annotated enhancers contained at least one hinge element (interquartile range 15.6% to 24.8%) (**Figure 5e&f, SI Figure 9**). In contrast, hinge positions were depleted of heterochromatin modifications with H3K9me3 occurring 0.13% of the time (interquartile range 0.1% to 0.17%) and H3K27me3 occurring 1.2% of the time (interquartile range of 0.9% to 1.8%, **Figure 5g&h, SI Figure 9**). Collectively this indicates, as proposed, that the act of transcription could facilitate the generation of domain geometry in a predictable manner.

### Genome geometry suggests transcriptional loops are efficiently packed

In sum, H1-H4 are supported by experimental evidence. While individually they can be explained by alternative mechanisms, the novel hypothesis that exons are non-randomly coupled to NE to generate domain volumes, in our view, best explains the sum of the evidence. Having demonstrated that the volumetric pattern of domains could be projected onto the positioning of exons, introns, and intergenic segments, we then studied whether they intersect with gene transcription. To do so, we tested if the proposed theory would indicate that transcriptionally active loops (mediated by Pol II), are packed in space. Two features we have proposed could be manifested in chromatin loops generated by Pol II^[62]^. The first is that some loops must represent durable domain volumes with compositions that mirror the ratios we described.

Durable structures could produce high-frequency loops within a population due to spatial confinement. Therefore, we used publicly available data through ENCODE to analyze the behavior of Pol II generated loops on ChiaPET^[62]^. We partitioned loops into very strong loops (>20 events) and compared them to more transient loops (<5 loop events). We then analyzed the composition of these two groups. Consistent with the novel hypothesis, very strong loops contain exon/NE ratios that are consistent with volumetric ratios of domains (**SI Figure 10**). Likewise, more transient loops spanned a spectrum of states including a mix of exon/NE segments, suggesting stretched as well as packing configurations (**SI Figure 10**).

To test if stable loops are a packed distribution of sizes, we utilized publicly available ATAC-Seq and ChIP-Seq. Similar to the geometry of domains guiding chromatin enzymes, the Tn5-transposase would interact with the outer zone based on the packing of the loop. If loops are instead a chain, then accessibility will be a linear function of loop length. Consistent with the packing properties proposed, accessibility is a power-law of length observed in strong loops (**SI Figure 10**). This occurs even upon depletion of RAD21, indicating that the generated volumes are not dependent on cohesin extrusion, which is consistent with findings on ChromSTEM imaging that mature domains remain even upon RAD21 depletion^[5,6]^ (**SI Figure 10**). Similarly, analysis of these volumes mirrored the proposed packing observed by segmenting the genome from the proposed hinges (**Figure 5c, SI Figure 10**). This is also observed experimentally (**Figure 5k, SI Figure 10**): the chromatin modifications are a power-law of non-exonic length within the loop segment. Features that represent the core of domains (H3K9me3, H3K27me3, **Figure 5k, SI Figure 10**) were at the proposed shorter depths (*p* ranging from 1 to 50 basepairs, **Figure 5f**) whereas Pol II and euchromatin aligned with predicted exon positions.

### Geometry suggests packing is associated with differentiation and oncogenic risk

We then explored the risks and benefits of the proposed geometric system. The process of domain formation and maturation echoes that of reinforcement learning systems in artificial intelligence and neural networks^[6,63,64]^. Inputs (signaling cascades, mechanical force) intersect with the current state (existing domains, ionic conditions, nuclear volume, and nucleosome remodeling complex concentration) that generate an output (RNA synthesis, domain activation/modification/degradation). Since the output modifies the state (domains), a future input experiences a different state that was modified by the prior inputs. By organizing the human genome into several thousand domains guided by transcriptional inputs determined by the existing state of the cell, cells can produce coordinated behaviors across a tissue. The domains encode a crucial feature for multicellular systems: a mechanism to produce coherent, reinforceable, and predictable memories of prior events. This represents an efficient, non-mutational system to increase complexity since geometry (packing) stores states. If this is true, genes active in terminally differentiated human tissues will be primarily power-law compositions, based on the need to maintain dynamic function for decades.

To test if this is indeed observed, we analyzed differentiated tissues from each germ layer: esophageal mucosa (endoderm), cardiac muscle (mesoderm), and cortical neurons (ectoderm). We utilized GTEx expression data from these three sites and selected genes that were preferentially associated with each tissue (**SI Table 1**)^[65]^. We observed conservation of the domain geometry system for genes involved in maintaining tissue function across the human lifespan (**Figure 6a&b**). Given this finding, we explored whether transcription factor families were similarly organized. Embryogenic development is a complex process with the timing of transcription factor activation impacting tissue formation. Therefore, we analyzed transcription factors in relation to developmental timing. We found that genes of early development (pluripotency factors, HOX genes)^[66,67]^ favored exon enrichment (linear structure or very small domains) whereas end-organ factors (e.g. MITF^[68]^ or RUNX2^[69]^) would generate large, power-law domains (**Figure 6c&d, SI Table 2**). However, by the proposed nature of a hinge guided by transcriptional activity, a portion of genes are placed in relatively risky conditions as they require being spanned between multiple reaction volumes. The risk to gene segments arises because they are exposed to environmental conditions and are at risk for entanglement events. As a result, our hypothesis predicts that these segments would be at a higher risk of mutations across all cancers. To test if this is the case, we investigated the mutation frequency of Tier-1 oncogenes compared to the likelihood that these genes overlap with a hinge position. We utilized the reported mutation frequencies across all tumor types generated by ROSETTA, which analyzed frequencies from publicly available cancer genomics sources (i.e. TCGA/TARGET Program)^[70,71]^. As these frequencies span across all tumor tissue types, they would provide an understanding of a conserved mechanism across cancers independent of tissue-specific factors. The null hypothesis is that no association would be observed and that mutations would be independent of the generated geometry. Remarkably, we observed that the frequency of oncogenic mutations is strongly correlated with the likelihood of a gene containing a hinge position (**Figure 6 e&f**). Indeed, many crucial Tier-1 genes (both oncogenes and tumor suppressors) overlap with the presence of a hinge including TP53, BRCA1/2, ATM, RB1, IDH1/2, and PIK3CA (**SI Table 3**). Collectively, these findings suggest that the proposed geometry is paired with a risk to genes that are required to create bifurcations between domains. While further work is necessary to understand if positioning is causal, it suggests that oncogenic mutation frequency may be linked to geometry.

**Figure 6.**
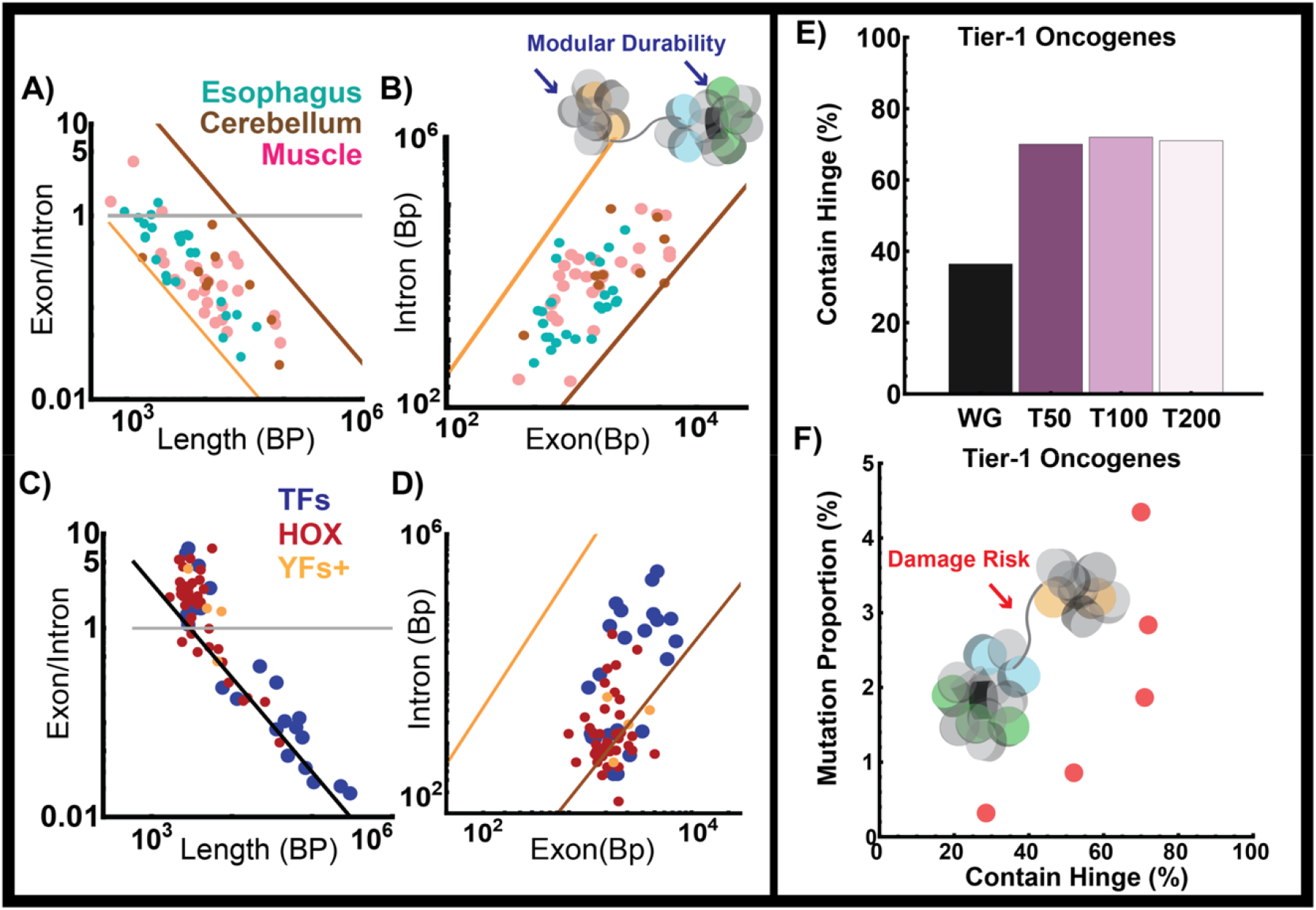
Power-law geometry produces a trade-off between modular durability and damage-risk. **A&B)** Analysis of geometric properties of genes that are primarily unique to the esophagus, cerebellum, and muscle tissue demonstrating power-law organization. **C&D**) Stem-cell related transcription factors (Yamanaka factors, YF) generally organize as linear geometries. Similarly, HOX genes demonstrate two phenotypes: a cluster with linear organization (values of E/I >1) and a group that organizes into power-law distribution. Transcription factors such as RUNX2 that define tissue function are primarily organized as power-law geometries. **E)** Analysis of the frequence of protein-coding genes containing a hinge position demonstrating that the 200 most frequent Tier-1 oncogenes contain at least one hinge position. WG – whole genome compared Tier-1 oncogenes by their frequency: T50 – top 50 genes, T100 – top 100 genes, T200 – top 200 genes. **F)** Analysis of Tier 1 oncogene frequencies demonstrates an acceleration then plateau in frequencies as the likelihood of containing a hinge increases. Genes that are less likely to contain a hinge element had a lower correlation with oncogenic mutation frequency. This effect appears to plateau at mutation frequencies occurring over 1% of the time.

### Power-law geometry parallels the emergence of organo-axial development

Since we propose that the benefit of the proposed geometry hypothesis is to produce durable cell states, we conclude by performing a comparative analysis across eukaryotic genomes to understand if it is a consequence of the act of splicing or an emergent regulatory mechanism. We specifically chose to analyze the following species due to their well-characterized genomes and existence of nucleosomes, introns, and splicing machinery: *S. cerevisiae* (a model of a monocellular organism with splicing and introns), *C. elegans* (a multi-cellular organism with simple organoaxial positioning), *D. melanogaster* and *D. rerio* (multi-cellular organisms with complex organoaxial positioning), and *M. musculus* (a non-primate mammal with complex organoaxial positioning)^[72]^. If this system is a byproduct of the act of splicing, all the investigated genomes would have similar geometries independent of complexity. Instead, if geometry built upon splicing to increase durable cell states, it would parallel the complexity of metazoan cell types. Consistent with the hypothesis that geometric encoding could facilitate organ complexity, power-law coupling of exons with introns appears to parallel organ specification **Figure 7a-h**). We observed the transformation from linear geometries with exon enrichment (*S. cerevisiae*) first toward a mix of power-law and linear structures (*C. elegans*) (**Figure 7a-h**). As complexity increased, there is a further transition from an equal mix of linear and small geometric assemblies to primarily geometric assemblies in *D. rerio* onwards. From the perspective of domain volume generation from the linear genomes, power-law geometry would require activation of one gene coming at the expense of a neighbor. Alternatively, transcription in linear genomes could occur in the span connecting domains, potentially indicating a mechanism by which the process progressed from a linear to a power-law assembly. While a linear assembly has benefits for rapid simultaneous synthesis, it is unlikely to efficiently produce memory and specialization. As a result of these considerations, it is worth noting that compaction within *S. cerevisiae* effectively translates into a 3-D barrier, whereas this is not the case in the assembly of domain volumes^[73]^. Instead, genomes built on a volumetric geometry can produce complex, highly dynamic, and durable states.

**Figure 7.**
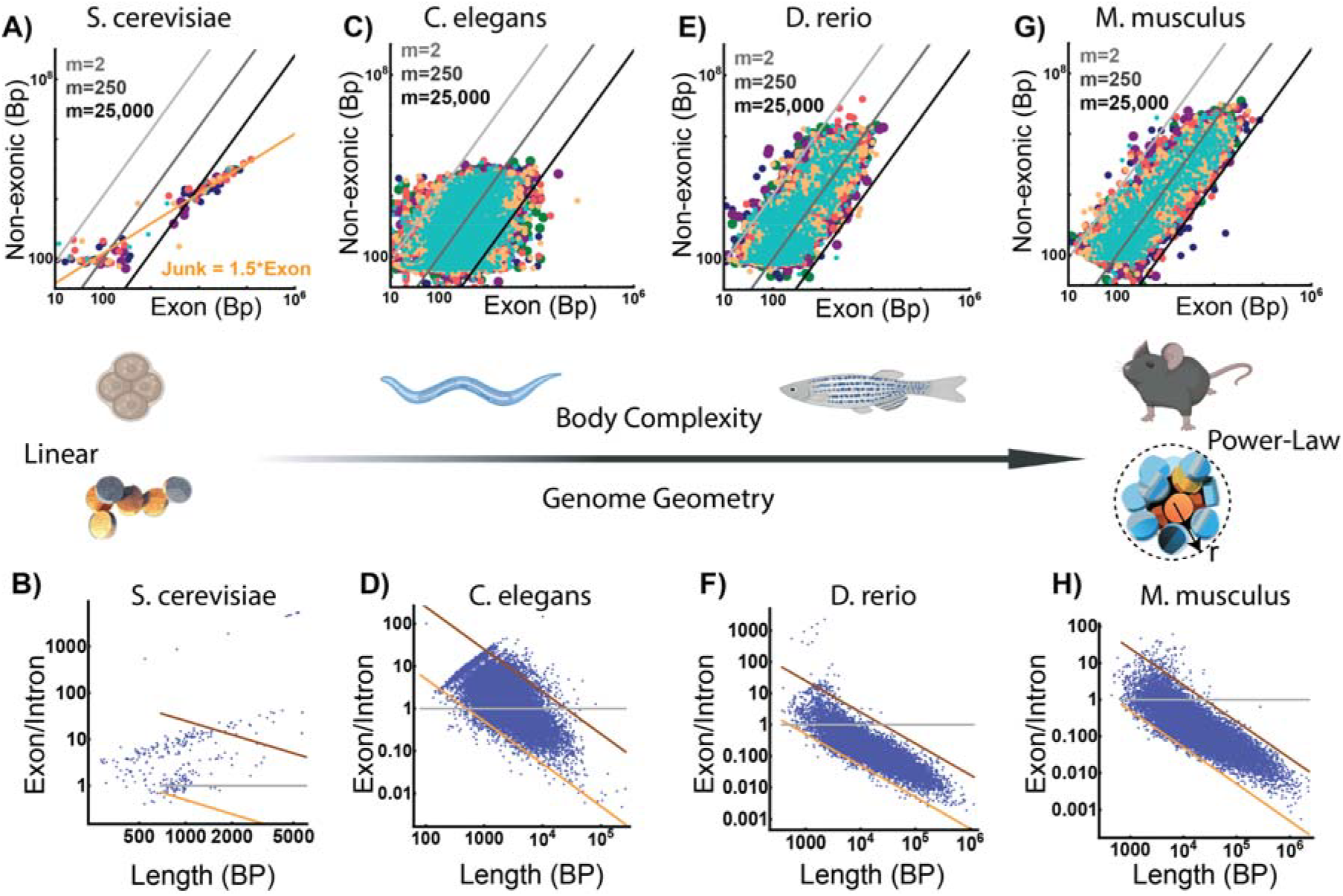
Packing geometry parallels body plan complexity in metazoans. **A&B)** Analysis of chromosomal architecture (**A**) and genes (**B**) demonstrates that the S. cerevisiae genome is most likely organized as linear beads on a string assembly. As gene length increases, the non-exonic content increases linearly to create a chain. **C&D**) Analysis of chromosomal (**C**) and genes (**D**) demonstrating a transition toward power-law assemblies. In C. elegans, genes appear to be equally split between linear assemblies (E/I >1) and power-law assemblies. **E-H**) We observe a transformation with increasing body-plan complexity in both genes and chromosomes of D. rerio (**E-F**) and M. musculus (**G-H**) that their organizational structure resembles the structure of genes and chromosomes observed in humans.

## Discussion

We set out to understand how packing domain information is stored within the genome in order to explain how a paradoxical decrease in accessibility of introns and intergenic segments is observed in transcriptional activation during muscle differentiation. We arrived at the novel hypothesis that the positioning of segments (exons, introns, intergenic) may encode 3-D packing information to generate nanoscale volumes. While additional investigation is needed to fully prove the implications of the proposed theory, it does introduce a mechanism for storing volumetric information for efficient transcriptional reactions (**Figure 1**)^[15,16,30,48,52,53]^. We first highlight that our findings demonstrate a crucial role for non-exonic DNA to act as ‘volumetric DNA’ in complex, multicellular eukaryotes to optimize chemical reactions involved in transcription^[42–44]^. This theory intriguingly suggests that the positional composition of human chromosomes may exist to produce efficient RNA synthesis (**Figure 2-5**). This is because while the vast array of the genome does not make RNA directly, length-based positioning would create efficient reaction volumes.

There are immense benefits from the proposed system: space and information optimization, durability, efficiency, simultaneous enzymatic processing, and geometric modulation to increase degrees of freedom (**Figure 1&2**).

Finally, it is notable that while most of the human, mouse, and zebrafish genomes appear to organize by a geometric principle (**Figures 7**), the transformation from linear, exon-rich configurations to power-law volumes correlates with the degree of body plan complexity. In future work investigating individual gene configurations, the framework demonstrated here could provide an understanding of how a beneficial epigenetic projection (3-D domain volumes) becomes encoded as a sequence-specific segment. This would require the ability to model and test specific gene configurations. Likewise, it generates questions as to whether DNA repair and replication enzymes behave like Pol II as a function of physiochemical conditions and the position of elements. We anticipate that future work can address these questions as the central hypothesis is mechanistically explored. These investigations could dissect how packing information is passed through the cell cycle, and how TADs and cohesin loops interact with this system. With those considerations in mind, we describe several transformative hypotheses to test across different disciplines based on the presented theory in addition to mechanistically testing the hypothesis proposed here:

### Hypothesis 1) Domain geometry facilitated the rapid emergence of body plan diversity

The transformation of chromosomes from linear to power-law assemblies parallels increasing body plan complexity (**Figure 7**). We naturally wonder if this converges at the time of the Cambrian Expansion^[74–76]^ as domain geometry can be a non-mutational system to increase degrees of freedom. This occurs because changing configurations depends on modifying spacing and position without requiring sequence mutations. This hypothesis can be tested by using the framework described here and then measuring the change in exon sequence composition compared to the change in arrangement. If evolutionary trajectories are found to be encoded in the arrangement of elements, it would demonstrate that selective pressure converges on gene position, orientation, and segment lengths to generate cohesive assemblies. What we perceive as ‘neutral drift’ from the sequence perspective is potentially balanced by selection for maintaining volumes. This allows increased sequence sampling of these regions mutationally for potentially beneficial states while maintaining a ‘default’ state as volumetric elements. Such findings would suggest that genomic selection is driven by the benefit of the system, and not necessarily the gene alone^[42,43]^.

### Hypothesis 2) Volume stabilization of mature domains defines cell response across decades

Nuclear swelling and heterochromatin loss are hallmarks of aging across human tissues^[53,77–79]^. Tissue development is defined by the non-random deposition of heterochromatin^[80]^. The volume ratios in packing domains reflect these states^[48,81,82]^. With this in mind, several diseases may be influenced by domain degradation over time. For example, among the risk factors for Alzheimers is the read-through fusion of ApoE and Tomm40^[83]^ (**Figure 1**). Could nuclear expansion and heterochromatin loss shift reaction volumes to produce transcriptional ‘fusions’ by positioning segments along the ideal zone as a single element^[84]^? Could a similar process be involved in inflammatory diseases since the misfolding of domains could create transcriptional memories that sustain inflammation^[85]^? Finally, in cancer, chromosome fusions, fragmentation, and copy number variations are significant prognostic factors. We observed that oncogene mutation frequencies were correlated with localization to a hinge segment (**Figure 6**); could these be non-random events that are related to how structure responds to local conditions? The nanoscale and microscale transformation of the nucleus is similarly a hallmark of malignancy and chemoresistance. Increasing the total genomic content, shifting the positioning of genes, and altering nuclear volumes generates unexpected geometries and the loss of coherent responses to the same stimuli. Based on the principles of domain geometry, one could prevent these events by targeting the physiochemical conditions that define the structure of domains^[86–88]^.

### Hypothesis 3) Domain geometry accommodates latency

Among the mysteries of transcriptional patterns is an unused surplus of binding sites for key transcription factors (e.g. Myod)^[89–91]^. While it has been suggested that these are errors, we can reconsider them in terms of information latency based on domains acting as a computational system. The default state generates volumes when unused, which remain as a reservoir for new domains if conditions change. In effect, the default positions needed for a human body plan are encoded in the measured positions of exonic/volumetric pairs with some degree of error to preserve a non-default adaptation strategy. If these adaptations improve fitness, they could be hard coded for the next progeny. Repositioning of these elements and the deposition of transcription factor binding sites based on geometry could provide insight into how sequence specificity intersects with geometric specificity.

### Hypothesis 4) Chromatin is a geometric computational system

By considering physical properties of the genome at the intersection with transcription reactions, these elements mirror aspects of learning, computation, and neural networks^[63,64]^. Some of the positional elements observed mirror elements of logic operators or computations, such as “AND” where genes in a paired segment become co-transcribed and “OR” states where one gene competes with another for volume. In this context, the predictable spacing between exons, introns, and intergenic segments requires inputs from the state of the cell. Alterations in mechanics^[30,92]^, ions^[88]^, redox state^[93,94]^, and nucleosome remodeling enzymes^[53]^ can provide this information. Domains would then act as geometric processors, with the structures formed representing the intersection of inputs and the current state. While this is more challenging to test, manipulating gene positions or controlling the order of signals could produce insights into this process. Finally, from the perspective of synthetic biology, controlling volume positioning during the generation of artificial chromosomes may guide the ability to generate complex traits

Collectively, this novel hypothesis requires additional mechanistic investigation for causality. The proposed system currently invites new avenues of exploration of genomics at the intersection of molecular, physical, chemical, and computational properties. Grounded in the intersection of these fields, exploration into the influence of domain volume assemblies on cellular fitness and organism evolution could serve as a framework for future studies in critical processes such as DNA replication, repair, and splicing.

## Materials and Methods

### Geometric positioning of exons and volumetric DNA in relation to packing domains

We derive scaling relationships to assess whether non-exonic (NE) DNA is coupled with exons to generate volume assemblies. Comparison between this analytical model and sequencing and ChromSTEM experimental data suggests that non-exonic segments likely correspond to volumetric organization surrounded by exons behaving as a “wavy” line on the reaction zone of the volume provided by the non-exonic elements, with domain fractal dimension *D* inversely related to *γ*.

Chromatin packing domains are nanoscopic, heterogeneous mass-fractal structures. The transformation of a chromatin chain into volume is defined by how the length of a segment of the chromatin polymer within a 3D domain, M, scales as a function of the radial distance of the volume containing the polymer, *r* by M∼r^*D*^, where *D* is the fractal dimension. The chromatin volume fraction, *ϕ*, at the radial distance *r* is defined by:

1)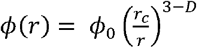 where *ϕ*_0_ is the chromatin volume fraction at *r*=0 (domain center) and *r*_*c*_ is the chromatin chain radius^[15,16,37]^. The total amount of chromatin in basepairs within the domain volume will therefore be:

2)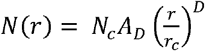

with *N* being the basepairs contained within the radius, N_c_ the number of basepairs within the chain and A_D_ the packing efficiency of the chain within a domain. Noting that A_D_ =1 indicates efficient packing throughout the domain volume.

For hard 3-D objects (e.g., a hard sphere), the number of basepairs, N_s_, within a zone shell of radius *Δ*R that is much smaller than the radius of the volume (*Δ*R << r) is:

3)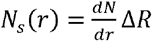

Substituting in Eq.(6), we therefore observe:

4)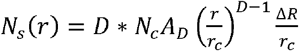

Accounting for the fact that the chain elements are a polymer that can go in and out of the hard shell (e.g., the “wiggly” line in **Figure 1**), it is instead necessary to take the fractional derivative to capture the behavior at the reaction zone:

5)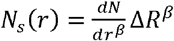

where *β* is the order (dimension) of the derivative and ranges from 0 ≤ *β* ≤1.

Utilizing the chain rule for a function f(x):

6)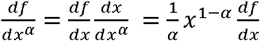

and Eqs.(9, 10), we therefore observe that the composition of the ideal zone is:

7)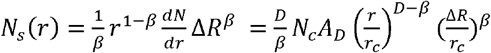

Assuming that the content of exons, E, in basepairs is approximately that of the contents of a domain ideal zone *E* ∝ *N*_*s*_ then the proportionality constant in this relationship is ∝ β. From this, we observe:

8)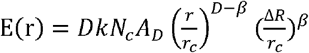

With *k* representing the fraction of the zone basepairs that are exons. Likewise, the length of a segment within a domain volume is the number of basepairs within that volume, *L*(r) = *N*(r). Transcriptional reactions occur at the ideal zone (the “Goldilocks zone”), the region where the balance between density stabilizes the intermediate complexes without overly limiting diffusivity of the reactant species^[87,88,95,96]^. For a domain limited by its ideal zone, the total length of the gene, L(r=R_gl_), and the exons, E(r= R_gl_) within domains to the goldilocks radius are:

9)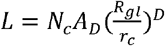

which indicates that

10)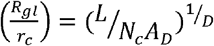

Solving for E using Eq.(12) utilizing the relation from Eq.(14) we can calculate the exon contents by

11)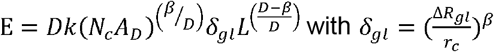

For simplicity, we can now let 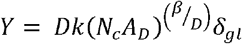 and c= 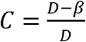 which simplifies Eq.(14) to

12) E= *YL*^c^ with *C* generally bounded between 2/3 and 1 due to the limits of *D* in cells of 2 to 3.

The existence of power-law scaling between exon length, E, and total gene length, L suggests a scaling relationship between the intron/intergenic segment and the exon. We observe an exponent, *γ* in Eq.(2) and Eq.(4) such that 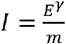.

If 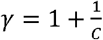then 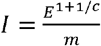, this would result in 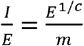 which by Eq.(15) becomes 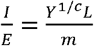. Given that Y is nearly 1, this results in 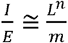 consistent with the experimental observations in Eq.(1).

Translating the power-law scaling described by *γ* into the mass-fractal dimension of domains, we observe that

13) 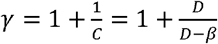as described in Eq.(4).

For a domain that is extends outside of its ideal zone, some NE regions may be found at r > R_gl_. Treatment similar to the one described above can be applied. From Eqs. 6 and 11, it follows that

14) 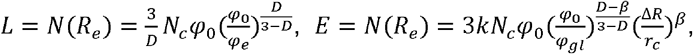,

where *φ*_e_ is the chromatin volume fraction at the radial distance corresponding to the outer bound of the domain, r=*R*_e_. In a special case of φ _0_ = φ_*gl*_, manipulation of equations 18 leads to

15) 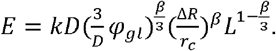.

Again, we see power-law scaling relationships among *l, E, l*:

16) 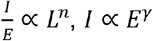, and *E* ∝ *L*^c^.

We now consider positioning of elements depending on the property of the exon segments in relation to the ideal zone elements defined by *β*. For *β*=1, the entire contents of the exon are within the hard shell. In contrast, for *β*=0, the exon portion is volumetrically distributed indicating that there is no geometry.

Indeed, in the most likely case, where exons constitute a portion of the ideal zone (a wiggly line) then 0 ≤ *β* ≤1 with *D* of domains ranging experimentally on ChromSTEM and in polymer modeling between 2 and 3 (2 ≤*D* ≤ 3)^[15,16,48]^, this results in 2 ≤ *γ* ≤3 observed experimentally.

When *γ*=2 and *β*=0, exons are distributed for any observed *D* (no geometric relationship). Conversely, when *γ*=3 and *β*=1, *D* =2 results in the limiting case of a chromatin polymer in a good solvent. The most frequently experimentally observed *γ* ∼ 2.3 is achieved when *β*∼0.7, indicating a substantial congruence between the ideal zone and exons, for experimentally (ChromSTEM) observed most probable *D* ∼ 2.8. It is worth noting that in this derivation the observed *γ* and *m* values plotted are those of the ensemble, and not the realized values for each individual gene. In essence, the entire genome is organized by this behavior, however, each individual gene may require segments from a neighboring segment in order to achieve the realized domain.

### Power-law genomic analysis

RefSeq genomes were obtained from Igv.org/app mapping to the UCSC genome browser assemblies. The selected genome assemblies were as follows: Human (GrCH38), Mouse (GRCm39), Zebrafish (GRCZ11), Drosophila melanogaster (dm6), and S. cerevisiae (sacCer3). These text files were converted to xlsx extensions and then imported into Mathematica v12 for subsequent analysis with custom-built code. Genes were then separated to analyze protein coding with assigned prefix “NM” and non-protein coding “NR”. The first isoform was selected for analysis of individual genes and compared to analysis with all isoforms. At the level of chromosome analysis, all isoforms were considered. Exon start and stop positions were used for segmentation and an reciprocal start/stop position for introns was generated. For simplicity, all exons were assumed to be part of the gene within the isoform variant. For chromosome wide analysis to account for the direction of transcriptional reading frames, genes were separated by their location to the positive or negative strand orientation for analysis. Multi-start or multi-stop exon overlapping events accounted for less than 7% of exon positions but were omitted in whole chromosome analysis for simplicity. Hinge elements were subsequently identified either for the whole chromosome in the read orientation (positive, negative) by mapping in the orientation of the read-frame (exon-NT) such that their size was below the predicted threshold: ∼200-300bp (1 nucleosome hinge) or ∼500-600bp (2.5 nucleosome hinge). The length of transcribed (exonic) and non-exonic (NE) sequences were summed in the intervening segments. The equations above were compared as described for the observed exon/intron ratio verses gene length, exon vs intron, and exonic vs NE in the respective figures. Randomization of only exons occurred by utilizing the generated lengths of exons and randomly repositioning these segments throughout the length of a chromosome. The remaining space was then defined as non-exonic. Randomization of an exon with its associated intron occurred by random permutation of the position of the shared elements along the generated lists for each chromosome.

Utilizing the Tissue Dashboard through the GTEx Portal, we subselected genes from the top 50 expressed within each respective tissues to represent different embryonic origins such that they generally did not overlap with other tissue beds. The selected tissues were the cortex (ectoderm), cardiac muscle (endoderm), and esophagus (endoderm).

### EU-RNA-Seq for Nascent RNA Analysis

#### Sample Generation

Nascent RNA was labelled, captured, and sequenced following the protocol reported by Palozola et al^[97]^. In brief, to selectively nascent RNA, HCT116 cells were treated with media containing 0.5 mM 5-ethynyluridine (EU) for 1 hour (Click-iT nascent RNA Capture Kit cat. no. c10365, Thermo Fisher Scientific). These RNAs were retrieved using Click-iT chemistry to bind biotin azide the ethylene group of EU-labeled RNA. The EU-labeled nascent RNA was purified using MyOne Streptavidin T1 magnetic beads. Captured EU-RNA attached on streptavidin beads was immediately subjected to on-bead sequencing library generation using the Universal Plus Total RNA-Seq with NuQuant® (Tecan) according to the manufacturer’s protocols with modifications. On-bead complementary DNA (cDNA) was synthesized by reverse transcriptase using random hexamer primers. The cDNA fragments were then blunt-ended through an end-repair reaction, followed by dA-tailing. Subsequently, specific double-stranded barcoded adapters were ligated and library amplification for 15 cycles was performed. PCR libraries were cleaned up, measured on an Agilent Bioanalyzer using the DNA1000 assay, pooled at equal concentrations and sequenced on a Novogene Nova Seq X with 50 BP paired end.

#### Data Analysis

Paired end nascent RNA transcripts were trimmed with TrimGalore v0.6.10 and aligned using Hisat2 v2.1.0. Alignment files were sorted, filtered, and converted to BAM format using Samtools v1.6. Coverage files were generated using Deeptools v3.1.1 and normalized using RPGC normalization. Finally, aligned filtered reads were counted using separate genome annotation files for intron, exon, gene bodies, and intergenic regions with Htseq v2.0.2. Gene body coverage plots were generated using the Superintronic package described in Lee et al^[98]^ by binning each gene body into 20 bins in the 5’->3’ orientation and calculating the average coverage in each bin for short, average, and long genes within introns and on exons.

### Estimation of compaction from nucleosome ‘beads on a string arrays’ compared to observed sizes

We consider the hypothesis that genes arrange as ‘open’ 10nm beads-on-a-string configurations to facilitate the RNA synthesis from a gene body in its entirety. One can calculate the length of the nucleosome array for thyroglobulin gene (TG) and Titan (TTN) compared to experimental observations as follows^[36]^. TG is ∼268,000bp and TTN is ∼304,000 which converts to an array of 1340 and 1520 nucleosomes for 200bp increments, respectively. Using the diameter of nucleosomes as 11nm, this produces chains that are 14.74 and 16.72 microns long for each gene. The reported median inter-flank distance in transcriptionally active state for each gene was observed as 703nm and 1104nm, respectively^[36]^. This 10-fold difference in length was accounted for by the introduction of stiffness, indicating the need for selective compaction of these genes even in their active state. A similar observation is observed within mouse neuronal cells in Rbfox1 (∼1.527Mbp in mice, ∼2.4Mbp in humans). In mice, Rbfox1 has a calculated chain length of ∼84 microns (200bp/bead) as ‘beads on a string’ but an experimentally observed transcription start site (TSS) to transcription end distance (TES) of ∼1 micron^[99]^.

This indicates an 84-fold compression between the TSS-TES during active RNA synthesis. A series of reaction volumes does not mean a gene does not undergo decompaction for transcriptional activation. Instead, it indicates decompaction likely transitions from large domains into a series of smaller domains. The smaller reaction volumes could be predictably encoded by the spatial positioning between exons, introns, and intergenic segments to create efficient reaction volumes.

### Chromatin Connectivity, Enhancer, Epigenetic Modifications and Tier-1 Oncogenes

The respective data of Chromatin interaction analysis with paired end tag (ChiaPET), Chromatin Immunoprecipitation with Sequencing (ChIP-Seq), and RNA-Sequencing Analysis (RNA-Seq) were all obtained from ENCODE^[7–9]^. The respective bed and bedpe files used were uploaded and listed in **SI Table 3**. The analysis for composition of ChiaPET loops were described as above where the total content in both orientations were considered for each chromosome. For analysis of element overlap with hinge positions, the identified 300bp hinges were used and the likelihood of any portion overlapping with a histone modification peak or enhancer peak was calculated. As a control variation in the read lengths covered by each element, the hinges observed when exons were randomly scrambled across the chromosome length were used. A two-tailed t-test comparing the observed frequency of overlap of the feature with a hinge per chromosome was compared to the observed frequency in the hinges of randomly generated segments.

RefSeq gene positions used to define the start/stop position of gene annotations as described in Almassalha et al^[53]^. Using the .bed files from ENCODE as above, we identified the mean location of each peak for the different marks with a p-value cut-off of at least <0.1. For each cell line, we then organized the Pol II-PS5 peaks by its density within each gene ranging from at least 1 to 10. We then calculated the distance of the Pol II-PS5 peak to the nearest peak of the respective histone mark. Using the cumulative density of distance, we calculated the median distance as a function of the number of Pol II-Ps5 peaks on a gene body. With respect to the per chromosome analysis, the total segment length of each mark was calculated for every somatic chromosome. We then normalized for the difference in chromosome length and plotted the coverage of Pol II-PS5 against each respective chromatin mark. In the case of human tissues, we instead plotted the association between heterochromatin and euchromatin since active RNA polymerase data was not available.

Analysis of oncogene mutational frequency compared to hinge positioning was performed by restricting the hinge length to 300bp or smaller (∼1-2 nucleosomes). The position of the hinges were mapped to protein coding genes. We then calculated the fraction of genes from a Tier-1 oncogene from the data within Mendiratta et al^[70]^. We then grouped the oncogenes into ascending groups and calculated the mean mutational frequency observed for each group. The values were then plotted compared to the observed frequency of hinges within genes within that group.

### Multi-color Single Molecule Localization Microscopy

#### Cell culture

HCT116 cells (ATCC, #CCL-247) were cultured in McCoy’s 5A Modified Medium (Thermo Fisher Scientific, #16600-082, Waltham, MA). The cell media were supplemented with 10% fetal bovine serum (FBS; Thermo Fisher Scientific, #16000-044, Waltham, MA) and 100 μg/ml penicillin-streptomycin (Thermo Fisher Scientific, #15140-122, Waltham, MA). Cells were cultured under standard conditions at 37°C in a humidified atmosphere with 5% CO2. Following detachment by trypsinization, cells were allowed to re-adhere and recover for at least 24 hours before further handling. Imaging was conducted when cell surface confluence ranged between 40–70%. For this study, cells were used between passages 5 and 20.

### Dual-color SMLM for EdU and histone modification

#### Cell Preparation and Fixation

After 48 hours from being seeded, the cells were incubated with EdU for 2 hours, followed by fixation for 10 minutes at room temperature with 4% paraformaldehyde in PBS. Fixed samples were washed three times in PBS for 5 minutes each. Secondary staining with AF647 via click-reaction chemistry was then performed according to the manufacturer’s protocol (ThermoFisher).

#### Permeabilization and Primary Antibody Staining

Samples were permeabilized and blocked using a buffer containing 3% bovine serum albumin (BSA) and 0.5% Triton X-100 in PBS for 1 hour. They were then incubated with rabbit anti-H3K9me3 or mouse anti-H3K27me3/H3k4me3 (Abcam) diluted in blocking buffer for 1–2 hours at room temperature on a shaker. Samples were washed three times in a washing buffer composed of 0.2% BSA and 0.1% Triton X-100 in PBS.

#### Secondary Antibody Staining and Imaging

The samples were incubated with goat anti-rabbit or goat anti-mouse AF488 (ThermoFisher) for 40–60 minutes at room temperature on a shaker. After incubation, samples were washed twice in PBS for 5 minutes each. Imaging was performed immediately, acquiring 10,000 frames per channel for each cell.

#### Dual-color SMLM for EdU and BrdU

After 48 hours of seeding, the cells were incubated with EdU and BrdU simultaneously for 2 hours. This was followed by fixation for 10 minutes at room temperature using 4% paraformaldehyde in PBS. The fixed samples were then washed three times with PBS, each wash lasting 5 minutes. BrdU and EdU staining was performed according the BrdU and EdU double staining protocol provided by Thermofisher.

#### SMLM reconstruction

The raw SMLM images were reconstructed using the built-in Thunder-STORM plugin in ImageJ. The camera setup parameters for the plugin were tailored to the imaging configuration. In our setup, we used a pixel size of 110 nm (calculated as the camera pixel pitch divided by the objective magnification), photoelectrons per A/D count of 1.09, and a base level of 0. The peak intensity threshold coefficient was adjusted based on the acquisition quality, typically ranging from 1 to 2.

#### Chromosome Paint

Human myoblasts were differentiated into myoblasts and chromosome painting was performed on myotubes as described previously^[29,39]^. Images were acquired on a Nikon Confocal Microscope or a Zeiss LSM 800 Confocal microscope. Imaging was done with a Plan Apo VC 100 × 1.4 NA oil objective as a multidimensional z-stack. The acquired 3D image stacks were then fed through imaging processing pipelines utilizing standard tools on Cell Profiler and Fiji pipelines performed the functions of translating images to maximal projections, calculating distance, object size, nuclear size, and radius^[100,101]^.

#### Declaration of generative AI and AI-assisted technologies in the writing process

During the preparation of this work the author(s) used ChatGPT in order to ensure readability. After using this tool/service, the author(s) reviewed and edited the content as needed and take(s) full responsibility for the content of the publication.

#### Data Sharing Plan

The generated analysis code and data will be made available upon reasonable request. Public sources of data are included in the summary tables for reference.

## Author Contributions

Conceptualization: LMA, KLM, MC, CD, IS, VB

Writing – original draft: LMA, KLM

Writing – review & editing: LMA, KLM, MC, CD, RG, JI, LMC, WSL, RN, PSD, IS, VB

Methodology: LMA, RG, IS, VB

Resources: LMA, KLM, PSD, IS, VB

Funding Acquisition: LMA, KLM, PSD, IS, VB

Supervision: IS, VB

Project Administration: IS, VB

Investigation: LMA, KLM, RG, JI, LMC, WSL

Validation: LMA

Formal Analysis: LMA, VB

Visualization: LMA, KLM

Software: LMA

Data Curation: LMA

## Acknowledgments

We appreciate the thoughtful discussion and feedback on this manuscript from Dr. Sui Huang. We appreciate the generous contributions from the funding sources listed below including the National Institutes of Health, National Science Foundation, Hyundai Hope on Wheels, Alex’s Lemonade Stand Foundation, Northwestern University, and Ann and Robert Lurie Children’s Hospital. Computational analysis of ATAC-Seq and EU-Seq data was supported in part through the computational resources and staff contributions provided by the Genomics Compute Cluster, which is jointly supported by the Feinberg School of Medicine, the Center for Genetic Medicine, and Feinberg’s Department of Biochemistry and Molecular Genetics, the Office of the Provost, the Office for Research, and Northwestern Information Technology. The Genomics Compute Cluster is part of Quest, Northwestern University’s high-performance computing facility, with the purpose to advance research in genomics. We appreciate the generous support from the ENCODE Consortium in the generation and dissemination of publicly available datasets. We specifically want to thank the labs of J Michael Cherry, Charles Lee, Richard Myers, Bradley Bernstein, Thomas Gingeras, Barbara Wold, Peggy Farnham, Michael Snyder, and John Stamatoyannopoulo for the generation and publication of the data utilized within this manuscript. We similarly appreciate the Atlas of Enhancers for their generation and dissemination of enhancer datasets.

## Funding

National Science Foundation grant EFMA-1830961 (MC, IS, VB)

National Science Foundation grant EFMA-1830969 (VB)

National Science Foundation grant CBET-2430743 (VB)

National Institutes of Health grant R01CA228272 (WSL, IS, VB)

National Institutes of Health grant U54 CA268084 (WSL, LMC, LMA, KLM, MC, IS, VB)

National Institutes of Health grant U54 CA261694 (LMC, VB)

National Institutes of Health grant U01DK134321 (PSD)

National Institutes of Health grant R01DK135620 (PSD)

NIH Training Grant T32AI083216 (LMA)

NIH Training Grant T32GM132605 (LMC, VB)

Hyundai Hope on Wheels Hope Scholar Grant (KLM)

CURE Childhood Cancer Early Investigator Grant (KLM)

Alex’s Lemonade Stand Foundation ‘A’ award grant (KLM) #23-28271

Ann and Robert H. Lurie Children’s Hospital of Chicago under the Molecular and Translational Cancer Biology Neighborhood (KLM)

National Institutes of Health National Center for Advancing Translational Sciences KL2TR001424 (KLM)

Northwestern University Starzl Scholar Award (LMA)

