## Supplementary material for "Geometrically encoded positioning of introns, intergenic segments, and exons in the human genome": SI File

**Supporting Material:**

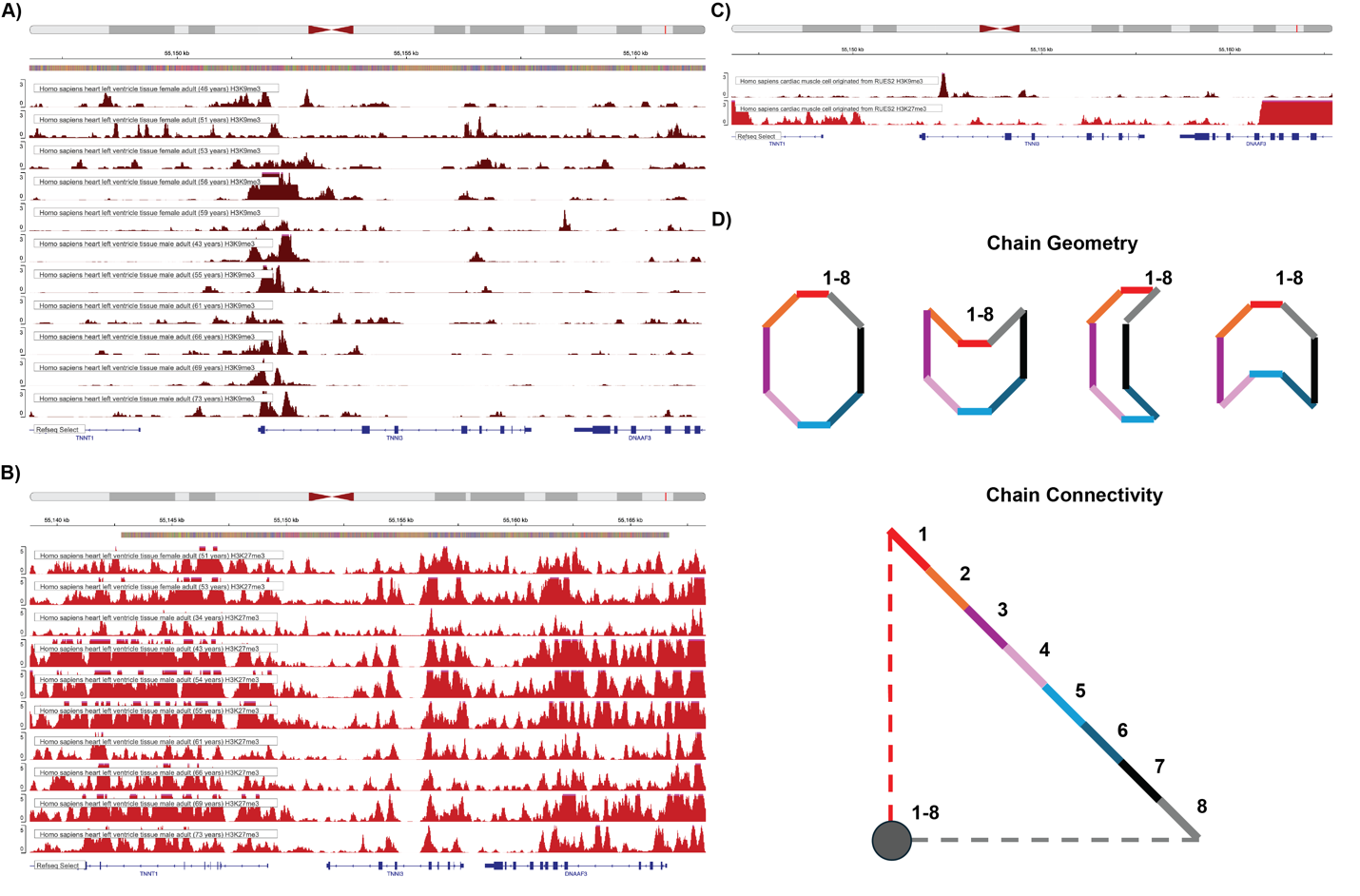
**SI Figure 1. Understanding genomic geometry is critical to understanding physiology. A**) Heterochromatin (H3K9me3) is deposited within the gene bodies of biallelic TNNI3 across multiple left ventricle samples independent of gender or cardiac functional status. Values scaled from 0 to 3. **B**) H3K27me3 is similarly deposited throughout intronic segments of left ventricle samples in TNNI3 independent of gender or cardiac functional status. Values scaled from 0 to 5. **C)** Deposition of H3K9me3 is observed within *in vitro* RUES differentiated cardiomyocytes, indicating that heterochromatin from left ventricular tissue is deposited in cardiac myocytes. Values scaled from 0 to 3. **D)** Schematic demonstrating that four different chain geometries each result in the same connectivity between distal loci (segment 1 is always connected to segment 8). See SI Table 5 for sample descriptions summarized from ENCODE.

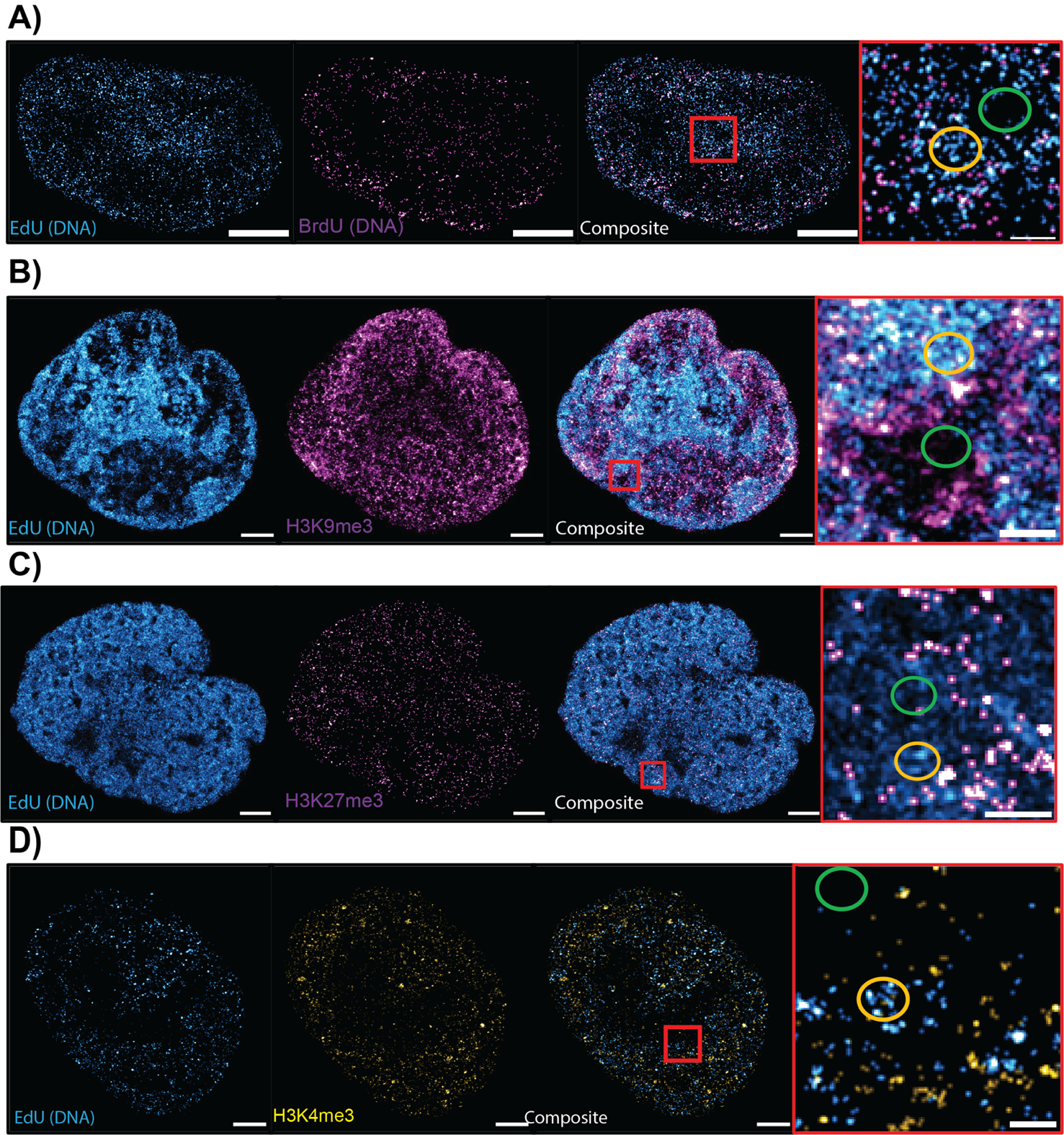

**Supplemental Figure 2) Steric exclusion of antibodies results in non-linear association between nucleosome modification and DNA density *in situ*. A)** Two color single-molecule localization microscopy of EdU and BrdU staining of DNA. EdU staining utilizing Click-it chemistry with a small molecule probe (~2nm) results in differential localization compared to antibody staining of DNA with BrdU (antibody ~10nm). Inset demonstrates staining behavior of antibody (purple) around the outside of a high-density domain (yellow) with comparable emission events to regions devoid of DNA (green) as identified based on EdU penetration. **B-C)** Two color single-molecule localization microscopy of EdU compared to H3K9me3 (**B**) and H3K27me3 (**C**) (both high-density heterochromatin) demonstrating localization *in situ* depends on size exclusion principles. H3K9me3 and H3K27me3 staining using antibody labeling results in accumulation near domain interior. **D)** Two color single-molecule localization microscopy of EdU and H3K4me3 (low density eu-chromatin) showing colocalization as a function of DNA density.

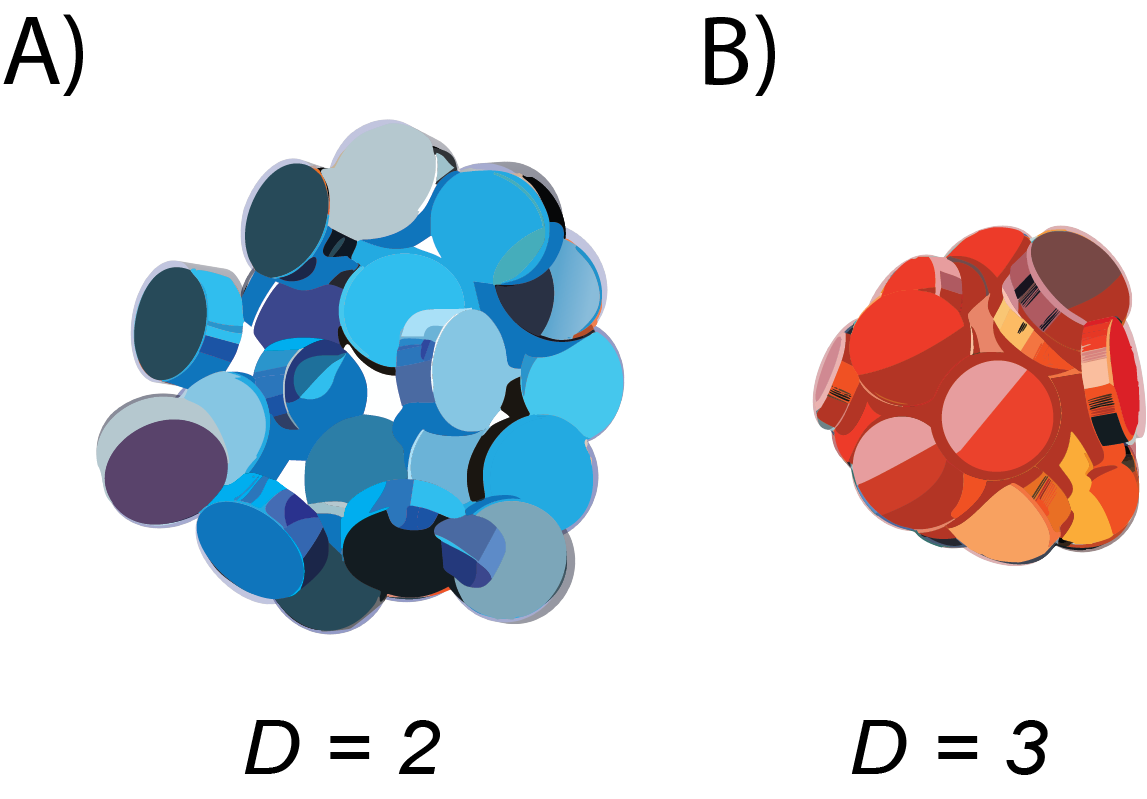

**SI Figure 3)** **Representation of the fractal polymer states.** **A)** Chromatin polymer as a homogenous structure with limited domain formation in the limiting case of *D = 2* **B)** Chromatin polymer as a confined, homogeneous media in the limiting case of *D = 3*. In both configurations, chromatin density is relatively uniform throughout.

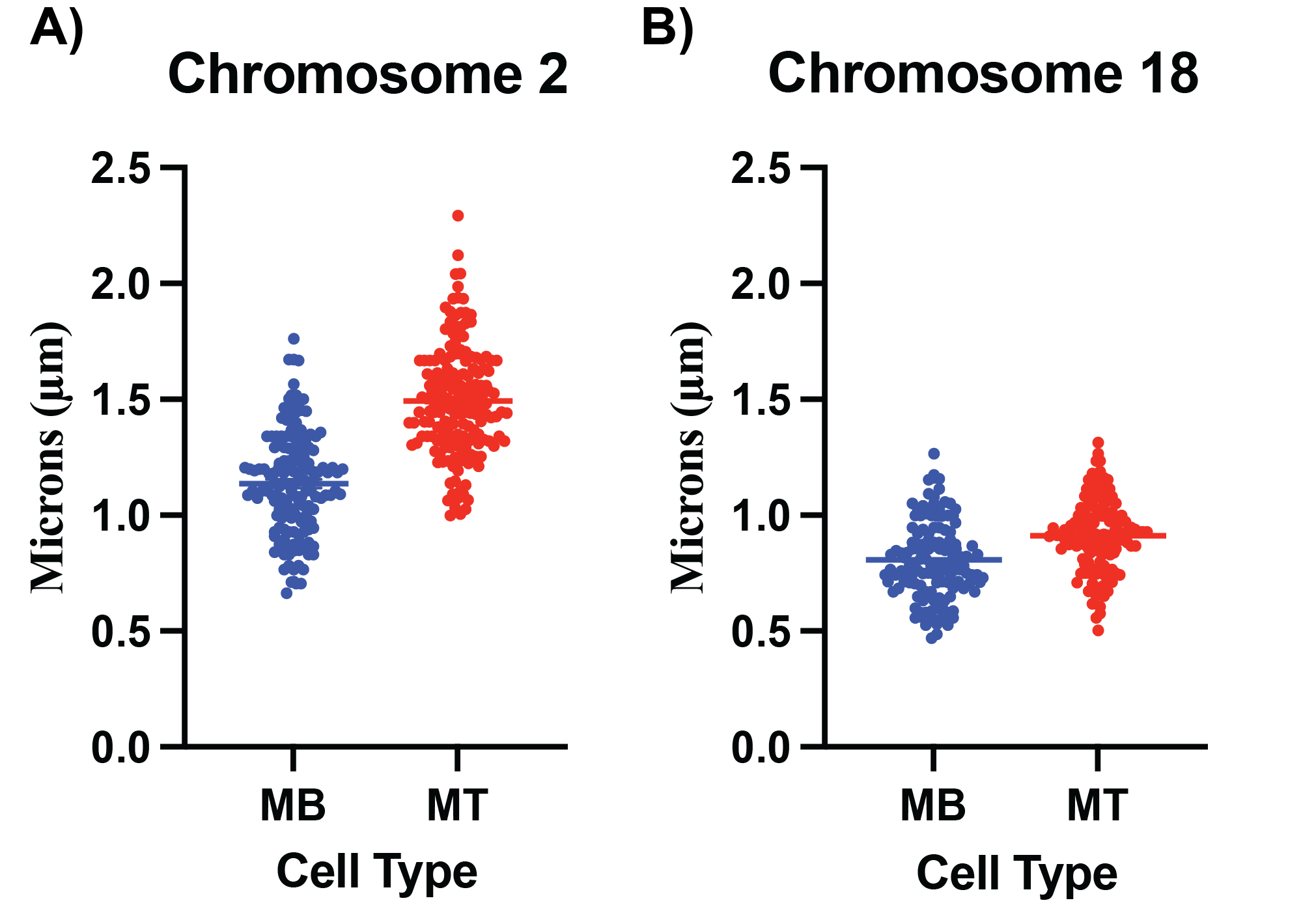
**Supplemental Figure 4) Measurement of chromosomal size in myogenesis. A)** Observed radius of chromosome 2 in immature muscle cells (myoblasts, MB) compared to mature muscle cells (myotubes, MT) demonstrating a radius of ~1-2microns in size for Chromosome 2 in mature muscle cells. **B)** Observed radius of chromosome 18 in immature muscle cells (myoblasts, MB) compared to mature muscle cells (myotubes, MT) demonstrating a radius of ~500nm-1.3microns in size for chromosome 18 in mature muscle cells. Even accounting for domain swelling due to FISH microscopy preparation, the occupied area for each chromosome would be comparable to a fully extended enhancer-promoter loop of 100kbp despite orders of magnitude differences in composition.

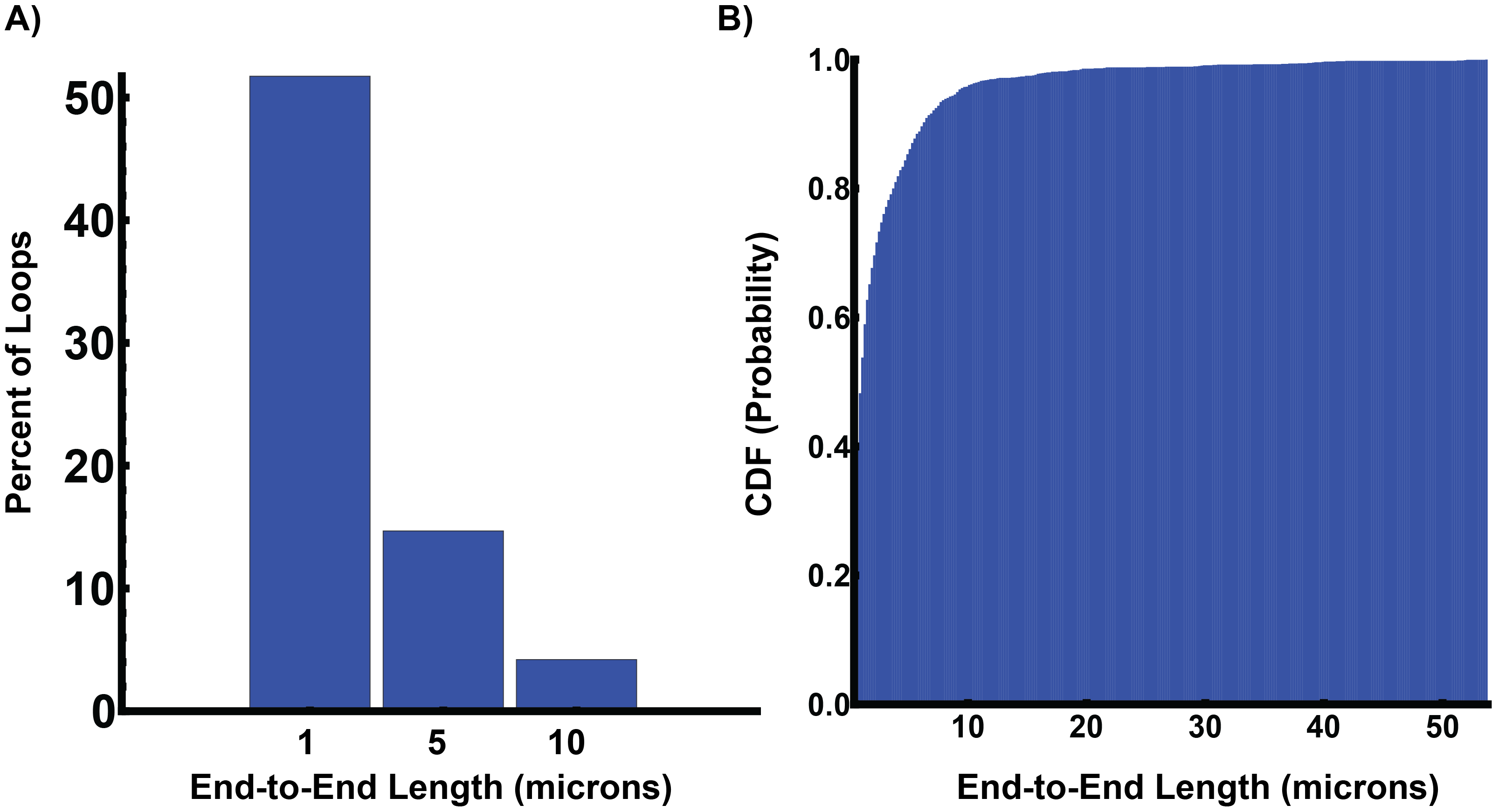

**Supplemental Figure 5) Analysis of loop sizes. A)** Quantitative estimates of the end-to-end length of observed loops generated by RNA polymerase II in HCT-116 cells demonstrating that 51% of observed loops would be approximately 1 micron long. Similarly, 14% of loops were at least 5 microns long and 4% were over 10 microns long. **B)** Cumulative density function histogram showing the distribution of loop sizes, with some loops extending to approximately 50 microns.

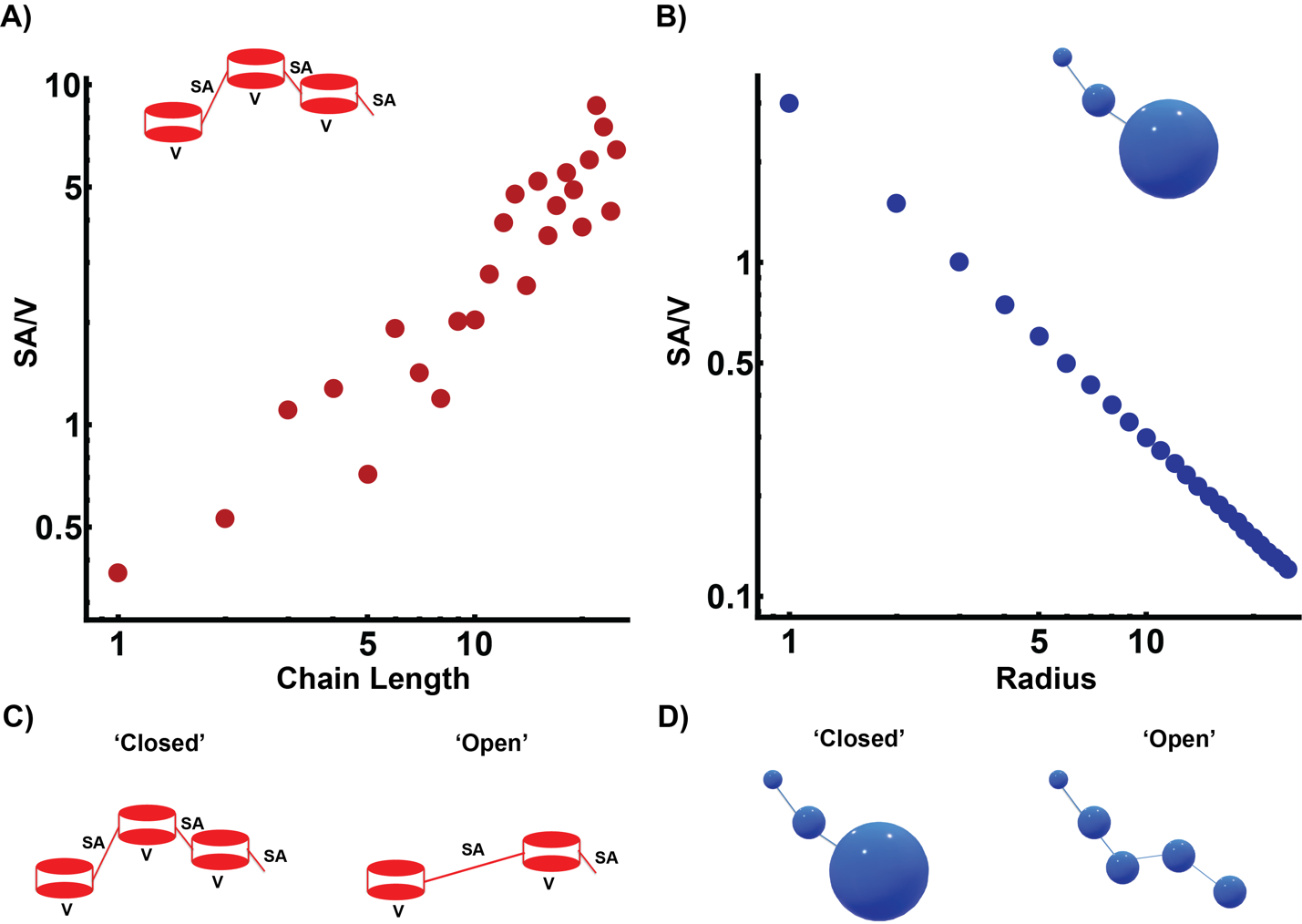

**SI Figure 6) Theoretical behavior of surface area to volume behaviors for chains compared to reaction volumes. A)** Beads-on-a-string configurations will have increasing linker segments as the chain increases in number. Linker DNA would represent the effective surface area of the chain whereas the DNA bound on a nucleosome could be considered the ‘volume’ component. Note that in this assembly, there is a linear increase in the SA/V of DNA content (linker/volume). **B)** Although chromatin is not organized into spheres, they are a useful approximation of the expected SA/V behavior. Organizing into reaction volume configurations will have an inverse relationship between the surface area to volume as the radius increases. **C&D)** Transitioning to an open configuration in a ‘beads on a string’ configuration results in increasing the linker distance without a nucleosome present. In contrast, increasing accessibility by generating reaction volumes would occur by creating a series of smaller volumes with a higher surface area to volume ration. In domain geometry, there would still be local compaction within each generated volume. The result would be an increase in accessibility that is a power-law of the radius instead of a linear increase in accessibility.

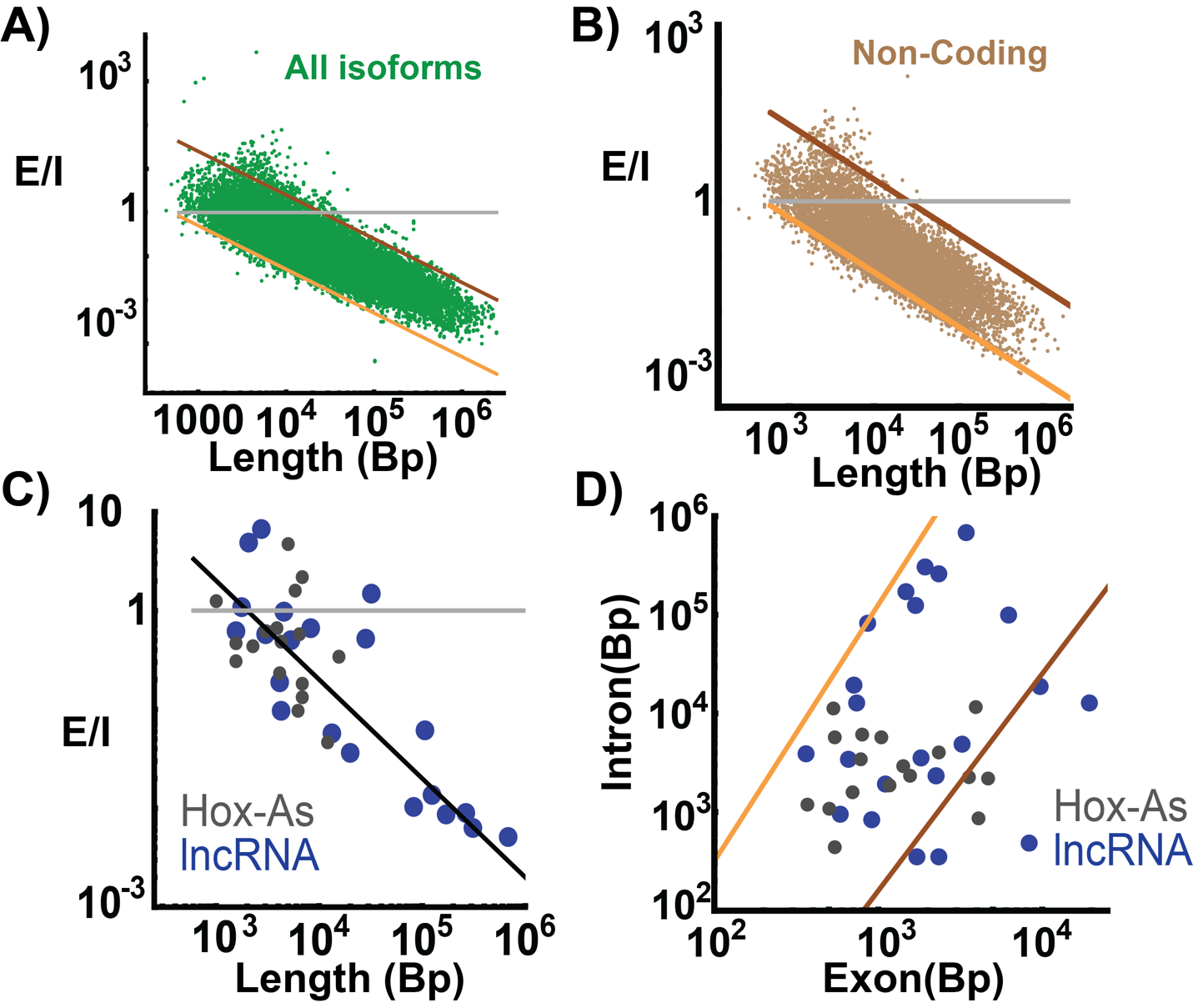

**SI Figure 7) Variant Isoform analysis and non-coding gene behavior. A)** Analysis of behavior of all protein coding genes demonstrating inverse relationship between surface area to volume ratio verses gene length across all isoforms. **B)** Analysis of behavior of non-protein coding genes (long non-coding RNA, pseudogenes, etc) demonstrating inverse relationship between surface area to volume ratio verses gene length independent of transcript products. **C-D)** Subselected long-noncoding RNA sequences (SI Table 3 for selected genes) demonstrates similar power-law behavior as that observed in protein coding genes independent of the RNA product function.

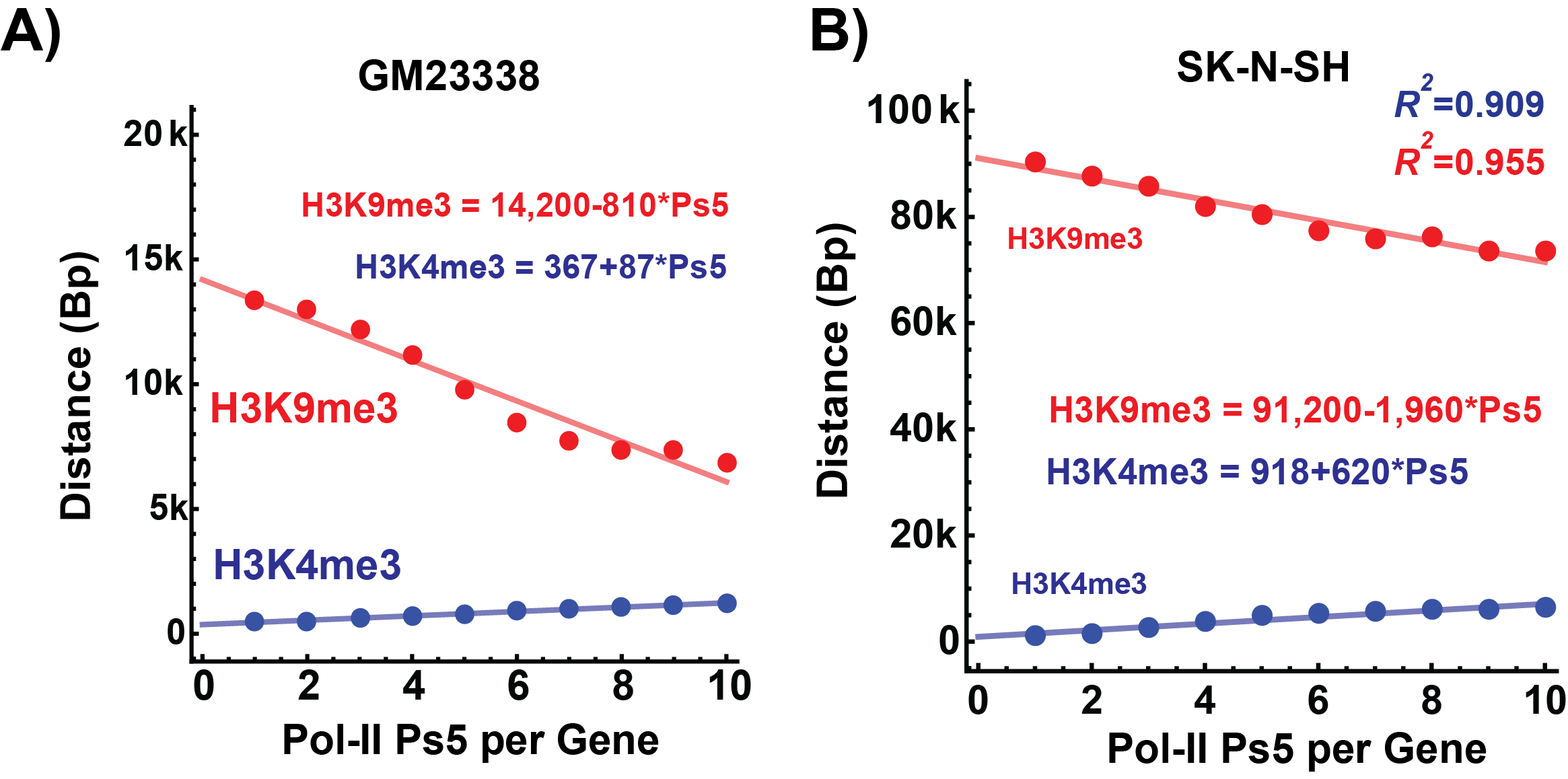

**SI Figure 8) Inverse distances in iPSC and SK Cells. A)** Distance relationship between active RNA polymerase (Pol II-Ps5) and the nearest H3K9me3 loci in GM23338 induced pluripotent stem cells (iPSC). The distance between active polymerase and the nearest core element, H3K9me3, is the shortest in iPSC cells consistent with cells enriched for small domain reaction volumes. Likewise, the distance decreases as the density of Pol II-Ps5 increases consistent with packing domain geometry. **B)** Distance relationship between Pol II-Ps5 and H3K9me3 loci in SK-N-SH neuroblastoma cell lines demonstrating a similar inverse relationship. Collectively across 4 cell line models (HCT-116 cells, Hep2G cells, GM23338, and SK-N-SH cells) an inverse relationship is observed consistent with the generation of packing domain reaction volumes.

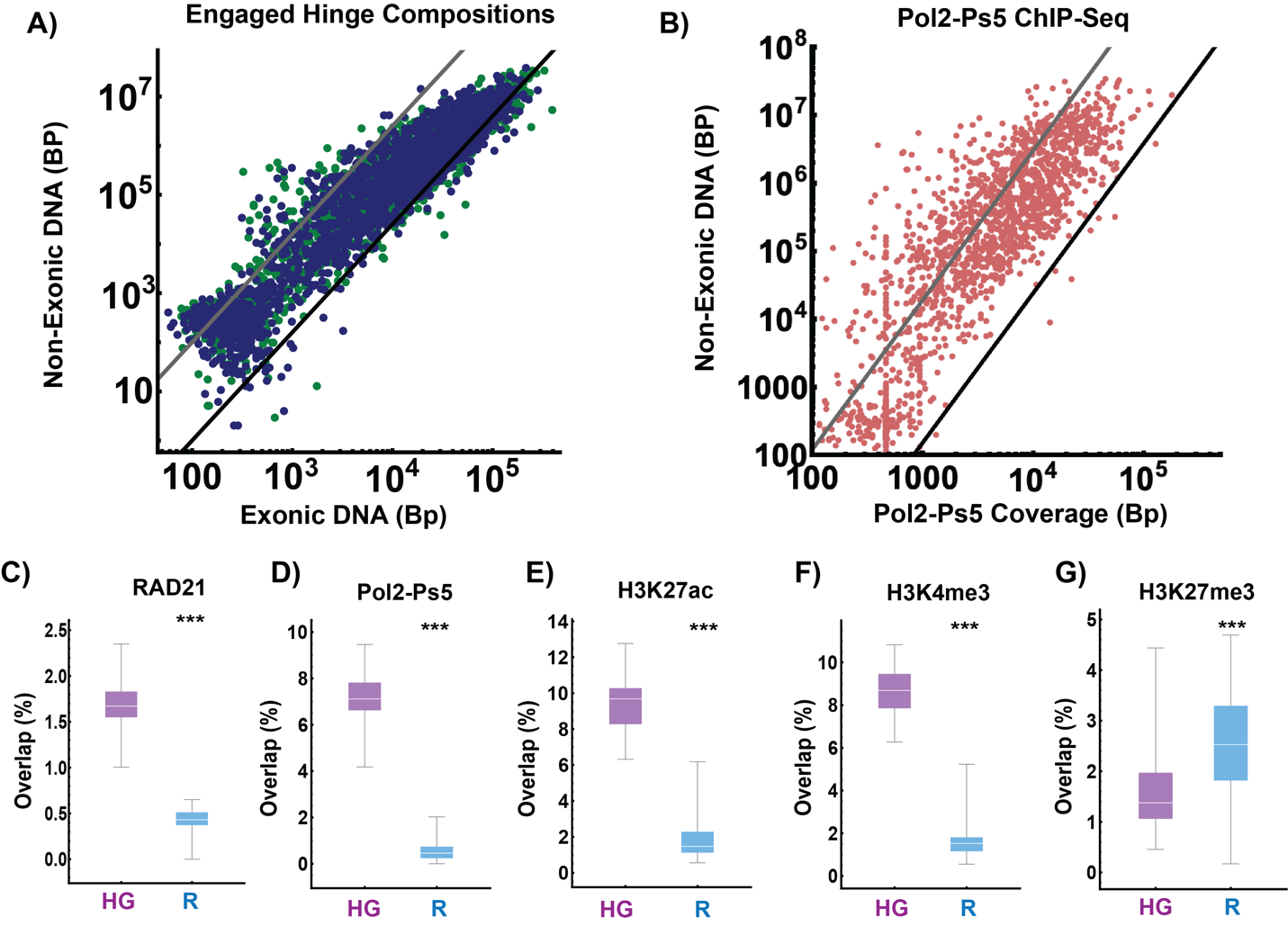
**SI Figure 9) Actively engaged hinge positions result in surface area to volume assemblies of exons and introns in HCT-116 cells. A)** Analysis of transcriptionally engaged hinge positions in positive (Green) and negative (Blue) orientations demonstrating surface area to volume compositions of exonic and non-exonic DNA. **B)** Analysis of RNA Pol II-Ps5 binding in HCT116 in engaged hinge segments demonstrating power-law coverage of Pol II Ps5 coinciding primarily with the position of exon elements. **C-G)** Analysis of ChIP-Seq binding frequencies of RAD21 (**C**), active Pol-II (**D**), and histone modifications (**E-G**) indicating hinges are preferentially associated with transcriptionally active features.

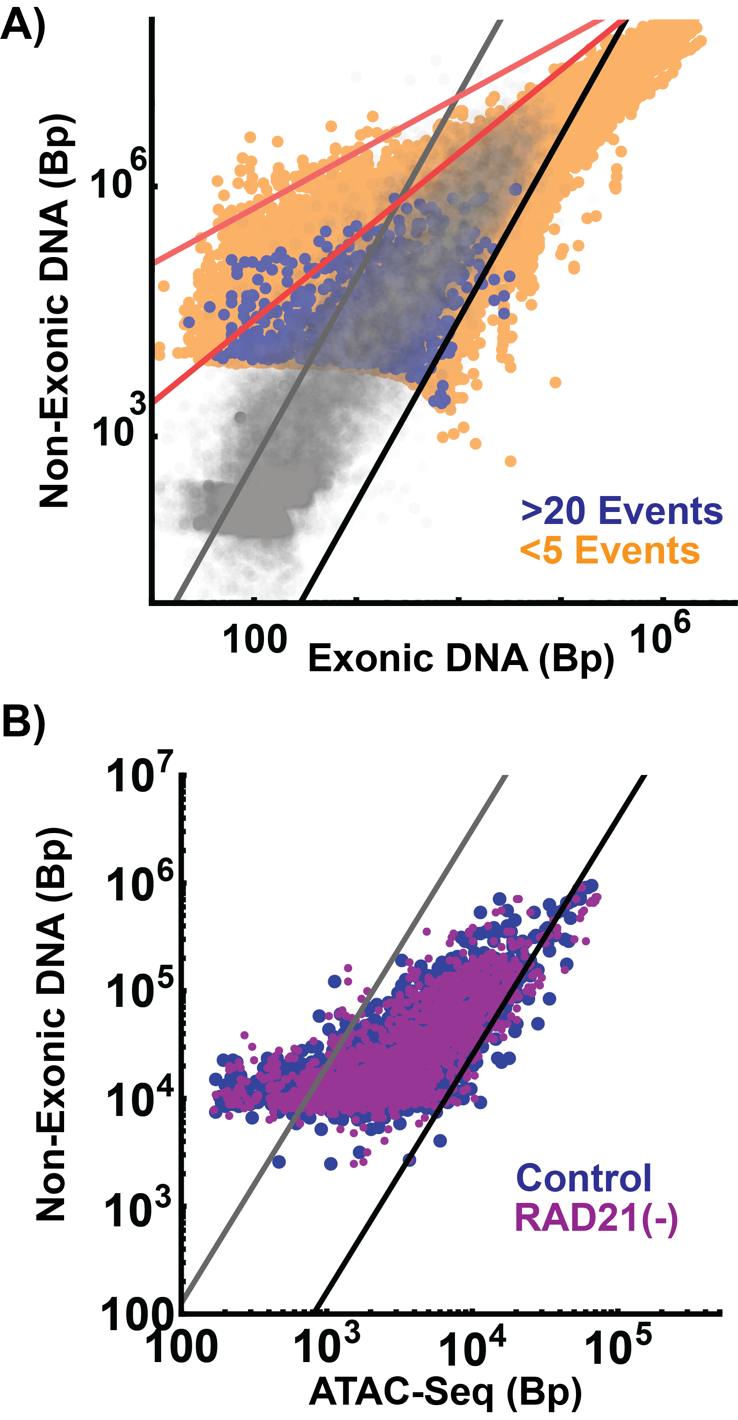

**SI Figure 10)** **RNA Polymerase loops organize into linear and power-law assemblies of exonic and non-exonic DNA. A)** Analysis of Pol-II mediated loops demonstrating that high frequency loops (>20 loop events observed) are composed as power-law assemblies of exonic and non-exonic DNA consistent with the generation of reaction volumes. **B)** Analysis of ATAC-Seq accessibility in HCT-116 high-frequency loops observed in (**A**) demonstrating that loop accessibility resembles a surface area to volume ratio as a function of the non-exonic length. This is consistent with high-frequency loops assembling into reaction volumes (packing domains) with non-exonic DNA predictably supplying the amount of volumetric DNA.

SI Table 1) Genes selected for each tissue

SI Table 2) Transcription factors, Yamanaka factors, and non-coding RNA genes selected for analysis

SI Table 3) ENCODE and Atlas of Enhancer accession information

SI Table 4) Tier 1 Oncogene hinge analysis

SI Table 5) Summary of patient information from ENCODE regarding cardiac functional status
